# Ferroptosis is a Physiologic Vulnerability of Iron-Recycling Macrophages

**DOI:** 10.64898/2026.06.22.732688

**Authors:** Miguel Mesquita, Maria Pires, Sara Violante, Jamil Zola Kitoko, Martim Costa, Sonia Trikha Rastogi, Rui Martins, Silvia Cardoso, Manuel Tanqueiro, Ana Figueiredo, Nelli Blank-Stein, Elvira Mass, Bindu Paul, Moises Mallo, Elisa Jentho, Miguel P. Soares

**Affiliations:** Gulbenkian Institute for Molecular Medicine (GIMM), Avenida Professor Egas Moniz – 1649-035 Lisboa, Portugal; Developmental Biology of the Immune System, Life & Medical Sciences (LIMES) Institute, University of Bonn; 53115 Bonn, Germany; Department of Physiology, Pharmacology & Therapeutics, Johns Hopkins University School of Medicine, Baltimore, MD 21205, USA; Department of Psychiatry and Behavioral Sciences, Johns Hopkins University School of Medicine, Baltimore, MD 21205, USA; The Solomon H. Snyder Department of Neuroscience, Johns Hopkins University School of Medicine, Baltimore, MD 21205, USA; Faculdade de Medicina da Universidade de Lisboa, Avenida Professor Egas Moniz 1649-035 Lisboa, Portugal

## Abstract

Iron deficiency anemia affects one-third of the global human population. Paradoxically, the daily iron required to fuel the production red blood cell (RBC) and prevent anemia is provided through its recycling from senescent RBC. This is achieved by splenic red pulp macrophages (RPM) that extract iron from the heme groups of hemoglobin (Hb). How these professional erythrophagocytic macrophages prevent intracellular iron flux from inducing cell death via ferroptosis is unknown. Here we show that SPI-C, the master transcriptional regulator of the erythrophagocytic lineage, orchestrates two redundant anti-ferroptosis pathways. One supports glutathione synthesis, via NF-E2-related factor 2 (NRF2), and the other relies on bilirubin production by biliverdin reductase A (BVRA). Genetic ablation of both pathways, but not either alone, sensitizes erythrophagocytic macrophages to ferroptosis, depletes RPM and increases the severity of iron deficiency anemia in mice. These findings reveal a central physiologic role of ferroptosis in the control of macrophage function, iron homeostasis and iron-deficiency anemia.

**Highlights:**

1. SPI-C enforces the antioxidant metabolic program of RPM.
2. SPI-C controls bilirubin production by biliverdin reductase A.
3. Bilirubin protects erythrophagocytic macrophages from ferroptosis.
4. Ferroptosis protection supports iron-recycling macrophages and limit iron deficiency anemia.

## Introduction

Iron is an essential micronutrient co-opted early in evolution^1^ to support reversible electron exchange across vital biochemical processes^2^. It is acquired from diet by enterocytes in the proximal small intestine^3^ and imported systemically via ferroportin 1 (FPN, *SLC40A1*)^4^. Iron is transported in plasma by transferrin^5^ and delivered to erythroblasts, through the transferrin receptor, supporting heme and hemoglobin biosynthesis during erythropoiesis. Perturbation of this inter-organ network, such as caused by iron deficiency, leads to anemia, affecting over one-third of the global population and a leading cause of human morbidity^6^.

Daily dietary iron covers roughly 3% of the erythropoietic demand^1,3^ and therefore iron is continuously recycled by splenic RPM^7–9^ that arise originally from yolk sac progenitors and are replenished throughout adult life by monocytes^10^. Erythrophagocytic macrophage differentiation occurs via a defined transcriptional program controlled by SPI-C, a heme-responsive member of the ETS (E26 transformation-specific) family of transcription factors^7,8^.

Constitutive expression of the heme catabolizing enzyme heme oxygenase 1 (HO-1, *HMOX1*) endows RPM with the capacity to extract iron from the heme groups of Hb^20^. Iron is exported from RPM via ferroportin-1 (FPN, *SLC40A1*) to support erythropoiesis^11,12^ while the open protoporphyrin ring biliverdin, is reduced into bilirubin by BVRA^13^.

The iron extracted from heme by erythrophagocytic macrophages poses a major challenge as it can catalyze lipid peroxidation, including that of polyunsaturated fatty acids (PUFA)^14–17^. If not controlled PUFA peroxidation ruptures phospholipid membranes^14,18^ triggering cell death via ferroptosis^14–17^. How RPM prevent iron from catalyzing PUFA peroxidation and causing ferroptosis is not understood.

Ferroptosis is regulated via glutathione-dependent and-independent pathways^19^. The glutathione-dependent axis is under the control of the transcription factor NRF2^20,21^, which regulates the expression of genes supporting glutathione synthesis^14^ to fuel glutathione peroxidase 4 (GPX4) activity and to limit PUFA peroxidation^22,23^. The glutathione-independent axis relies on the production of lipophilic radical-trapping antioxidants (RTA) that inhibit PUFA peroxidation^14,18,19,24,25^. These include ubiquinol (CoQ_10_H), produced via the reduction of ubiquinone (CoQ_10_), catalyzed by the NAD(P)H-dependent oxidoreductase ferroptosis suppressor protein 1 (FSP1/AIFM2)^24,25^.

RPM express constitutively high levels of BVRA, a NAD(P)H-dependent oxidoreductase that catalyzes the reduction of biliverdin into the lipophilic RTA^6^ and superoxide scavenger^2–4^ bilirubin^1^. While BVRA confers resistance to oxidative stress^26^, whether bilirubin is protective against ferroptosis is not known.

Here, we identify SPI-C, the master transcriptional regulator of the RPM lineage, as the orchestrator of a glutathione-dependent and-independent axis that protects erythrophagocytic macrophages from ferroptosis. The glutathione-dependent axis supports glutathione production via activation of NRF2 and the glutathione-independent axis relies on the production of bilirubin by BVRA. Together, these redundant cytoprotective pathways decouple erythrophagocytosis from ferroptosis, establishing ferroptosis as a major physiologic vulnerability of iron-recycling erythrophagocytic macrophages, including but not restricted to RPM. This integrated cytoprotective response is essential to support iron recycling and limit the severity of iron deficiency anemia, revealing ferroptosis as a therapeutic target against this global cause of human morbidity^6^.

## Results

### SPI-C controls the antioxidant transcriptional response to heme

(*Fig. 1; Fig. S1*) Erythrophagocytic macrophages are exposed to high levels of exogenous heme derived from Hb, mimicked experimentally by exposing bone marrow-derived macrophages (BMDM) to exogenous heme^7,8^. In response, BMDM activated a transcriptional program characterized by the up-regulation of 1,267 genes (*Fig. 1A-C*), including a cluster of 33 genes regulated by the transcription factor NRF2 (e.g., *Nqo1, Sod1, Txnrd1*). Other heme-responsive genes were involved in glutathione metabolism (16 genes; e.g., *Slc7a11, Gsta3, Gclm*), NADPH generation (4 genes; *Me1, Pgd, G6pdx, Idh1*), iron homeostasis (3 genes; *Slc40a1, Cp, Fth1*), heme/metal binding (2 genes; *Slc48a1, Alas1*), inflammatory (7 genes; e.g., *Ptgs2, Il1b, Il1a*) and immune (2 genes; *Tnf, Irf1*) responses as well as genes underlying chemokine activity (10 genes; e.g., *Cxcl3, Cxcl1, Cxcl2*). Conversely, BMDM repressed 1,260 genes in response to heme, including cell cycle regulatory genes (23 genes; e.g., *Ccnb1, Ccnb2, Cdk4*), genes involved in DNA replication (10 genes; e.g., *Mcm3, Mcm5, Rfc2*) and mitotic checkpoint genes (7 genes; e.g., *Cenpa, Bub1, Nek2*) (*Fig. 1A-C*). Of note, heme induced a coordinated regulation of biliverdin reductase A and B, whereby *Blvrb* was induced while *Blvra* was repressed (*Fig. 1A-C*).

**Figure 1.**
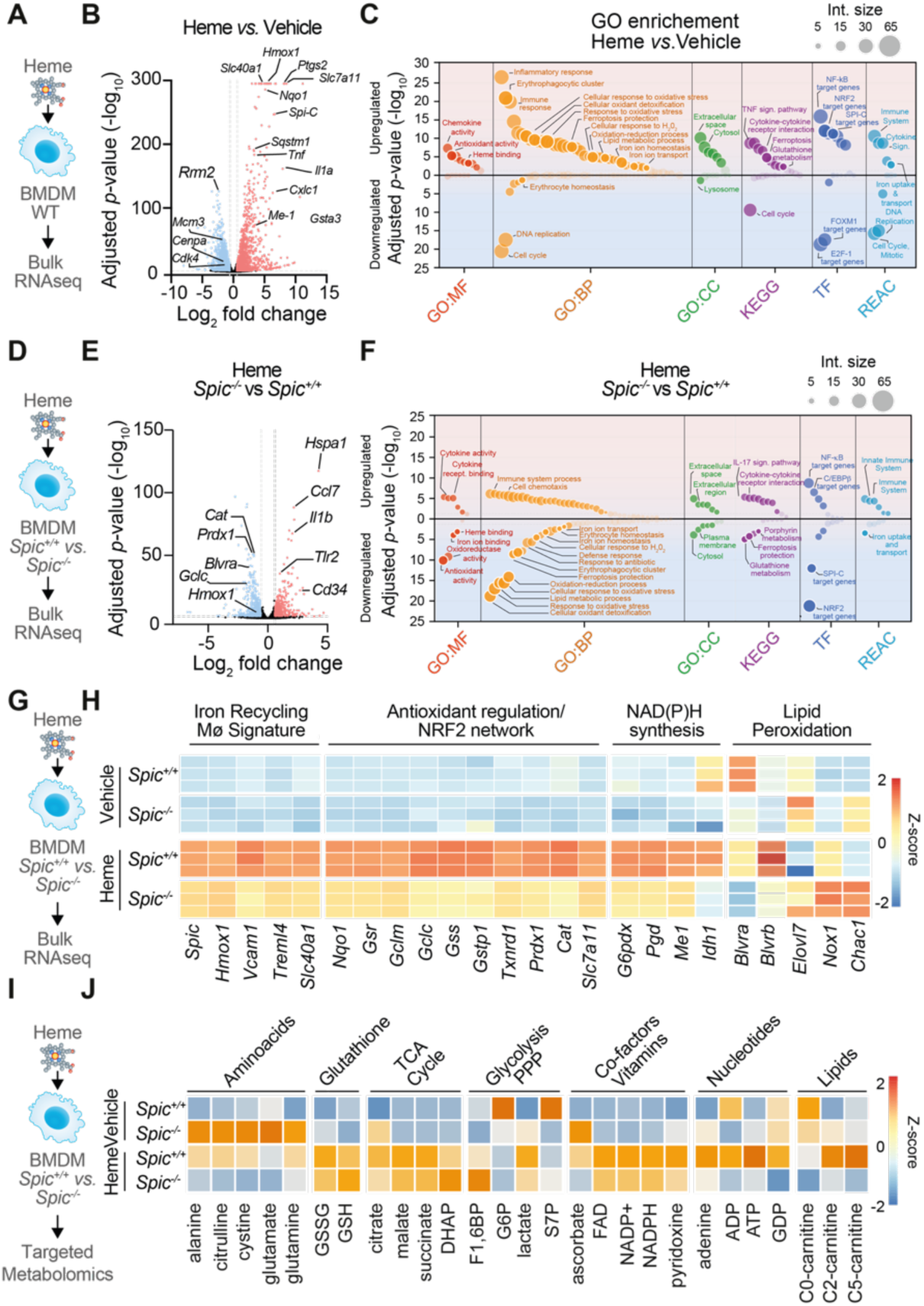
Heme drives SPI-C-dependent erythrophagocytic, anti-ferroptotic transcriptional and metabolic programs in macrophages. (A) Schematic representation of bulk RNA-seq analyzes of BMDM from C57BL/6J (WT) exposed to heme *vs.* vehicle, (B) Volcano plot of differentially expressed genes. Heme-upregulated and downregulated genes indicated in red and blue, respectively. Dashed lines denote significance thresholds (log₂ fold change and adjusted *p*-value). (C) Functional enrichment analysis of heme-upregulated and-downregulated gene sets across GO:MF, GO:BP, GO:CC, KEGG, TF, and Reactome (REAC) categories. Circle size represents gene set size; position along the y-axis reflects statistical significance. (D) Schematic representation of bulk RNA-seq analysis *Spic^+/+^* (WT) and *Spic^-/-^* BMDM exposed to heme *vs.* vehicle. (E) Volcano plot of differentially expressed genes of *Spic^-/-^vs. Spic^+/+^*. (F) Functional enrichment analysis of SPI-C-dependent upregulated and downregulated gene sets across GO:MF, GO:BP, GO:CC, KEGG, TF, and Reactome categories. (G; H Schematic representation of experimental procedure and heatmap of selected differentially expressed genes grouped by functional category. Same samples as in (E,F) (I, J) Schematic representation of *Spic^+/+^*(WT) and *Spic^-/-^* BMDM treated with heme *vs.* vehicle and subjected to semi-targeted metabolomics analysis. (J) Heatmap of selected metabolites grouped by pathway. RNA-seq data in (B, C, E, F, H) from one experiment with n=3 replicates *per* condition. Metabolomics data in (J) is from one independent experiments with n=6 replicates *per* condition.

To determine the relative contribution of SPI-C to the heme-responsive transcriptional program we generated *Spic-*deficient (*Spic*^-/-^) C57BL/6J mice, carrying a disrupting of the *Spic* locus between exons 3 and 4 (*Fig. S1A-C*). BMDM from *Spic*^-/-^ mice failed to express *Spic* mRNA (*Fig. S1D*) and presented the expected age-dependent depletion of RPM (*Fig. S1E,F*)^7,8^.

When exposed to exogenous heme, *Spic^-/-^* BMDM showed attenuated induction of NRF2-regulated genes (e.g., *Hmox1, Nqo1, Slc7a11, Gclc, Gclm, Gsr*) as well as genes involved in NADPH synthesis (e.g., *Me1, Pgd, G6pdx*) and iron handling (e.g., *Slc40a1, Fth1*), failing to execute the biliverdin reductase A and B switch, compared to the robust activation of these modules in control *Spic^+/+^* BMDM (*Fig. 1D-H, Fig. S1G*). In contrast, the repression of cell cycle and proliferative genes (e.g., *Ccnb1, Mcm3, Cenpa*) was maintained in *Spic*^-/-^ BMDM exposed to heme.

These data reveal that SPI-C controls a coordinated transcriptional response to heme encompassing the activation of a NRF2-dependent antioxidant defense, NADPH generation, glutathione metabolism and reprogramming of bilirubin production, while proliferative arrest occurs independently of SPI-C.

### SPI-C regulates the antioxidant metabolic response to heme (*Fig. 1; Fig. S2*)

To further establish the impact of SPI-C on glutathione, NADPH and other components of macrophage metabolism we performed targeted metabolomics from *Spic*^⁻/⁻^ *vs. Spic*^⁺/⁺^ BMDM. At steady state, *Spic*^⁻/⁻^ BMDM presented reduced glutathione (GSH) with a concomitant increase in oxidized glutathione (GSSG), collapsing the GSH/GSSG ratio, compared to control *Spic*^⁺/⁺^ BMDM (*Fig. 1I,J, Fig S2A-B*). Intracellular levels of TCA cycle intermediates were comparable between genotypes at baseline, with the exception of α-ketoglutarate (*Fig. 1I,J, Fig S2C-E*). *Spic*^⁻/⁻^ BMDM presented higher levels of arginine, citrulline, alanine, cystine, methionine and glutamine, compared to *Spic*⁺^/^⁺ controls (*Fig. 1I,J, Fig S2E*). This suggests that SPI-C regulates amino acid and glutathione metabolism while exerting a negligible effect on the TCA cycle at steady state.

The metabolic response to exogenous heme diverged between *Spic*^⁻/⁻^ and *Spic*^⁺/⁺^ BMDM (*Fig. 1I,J, Fig S2*). GSH was modestly higher in heme-exposed *Spic*^⁻/⁻^ *vs. Spic*⁺^/^⁺ BMDM, yet NADPH was reduced with a concomitant elevation of GSSG, collapsing the GSH/GSSG ratio (*Fig. 1I,J, Fig S2A,B*). Cystine, methionine, and glutamine, were all elevated at baseline in *Spic*^⁻/⁻^ BMDM, but conversely depleted upon heme challenge in *Spic*^⁻/⁻^ but not in *Spic*^⁺/⁺^ BMDM (*Fig. 1I,J, Fig S2C*). This is consistent with a failure to sustain precursor flow under increased oxidative demand, pointing to a defect in NADPH-dependent glutathione recycling rather than glutathione synthesis.

FAD, NAD⁺ and acetyl-CoA increased upon heme exposure in BMDM from both genotypes, indicating that metabolic activation downstream of heme catabolism proceeds independently of SPI-C (*Fig. 1I,J, Fig SF*). Within the purine nucleotide pool, ADP and GDP remained stable, while ATP increase upon heme exposure in *Spic*⁺^/^⁺ but not in *Spic*⁻^/^⁻ BMDM (*Fig. 1I,J, Fig S2G*), consistent with reduced energetic output despite preserved upstream cofactor responses. Acylcarnitine species diverged in parallel: free carnitine (C0-carnitine) declined more markedly in *Spic*⁻^/^⁻ under heme, while C2-and C5-carnitines rose only in *Spic*⁺^/^⁺ BMDM (*Fig. 1I,J, Fig S2H*), consistent with altered mitochondrial fatty acid handling.

The metabolic deficits of *Spic*⁻^/^⁻ *vs. Spic*⁺^/^⁺ BMDM suggests that SPI-C coordinates a transcriptional response to heme that promotes cystine import, glutathione recycling, and NADPH availability. This suggest that the transcriptional program regulated by SPI-C is essential to maintain and sustain substrate flow in response to exogenous heme when macrophages engage into erythrophagocytosis. How this transcriptional-regulated metabolic response impacts on ferroptosis is addressed below.

### NRF2 and BVRA provide an integrated cytoprotective response against ferroptosis (*Fig. 2; Fig. S3,4,5*)

To test the contribution of NRF2 to the cytoprotective response of macrophage against ferroptosis we generated *Nrf2^LysMΔ/Δ^* mice, carrying an *Nrf2* deletion specifically in myeloid cells (*Fig S3A-E*). Intracellular iron overload, induced by ferric ammonium citrate (FAC) failed to induce ferroptosis in *Nrf2^LysMΔ/Δ^*BMDM, similar to control *Nrf2^fl/fl^* BMDM (*Fig S3F,G*). Pharmacologic inhibition of GPX4 (1S, 3R)-RSL3 (RSL3)^5,6,7^ also failed to induce ferroptosis in *Nrf2^LysMΔ/Δ^* BMDM, similar to control *Nrf2^fl/fl^* BMDM (*Fig S3H,I*). This suggests that macrophages deploy an NRF2-independent axis to repress ferroptosis.

**Figure 2.**
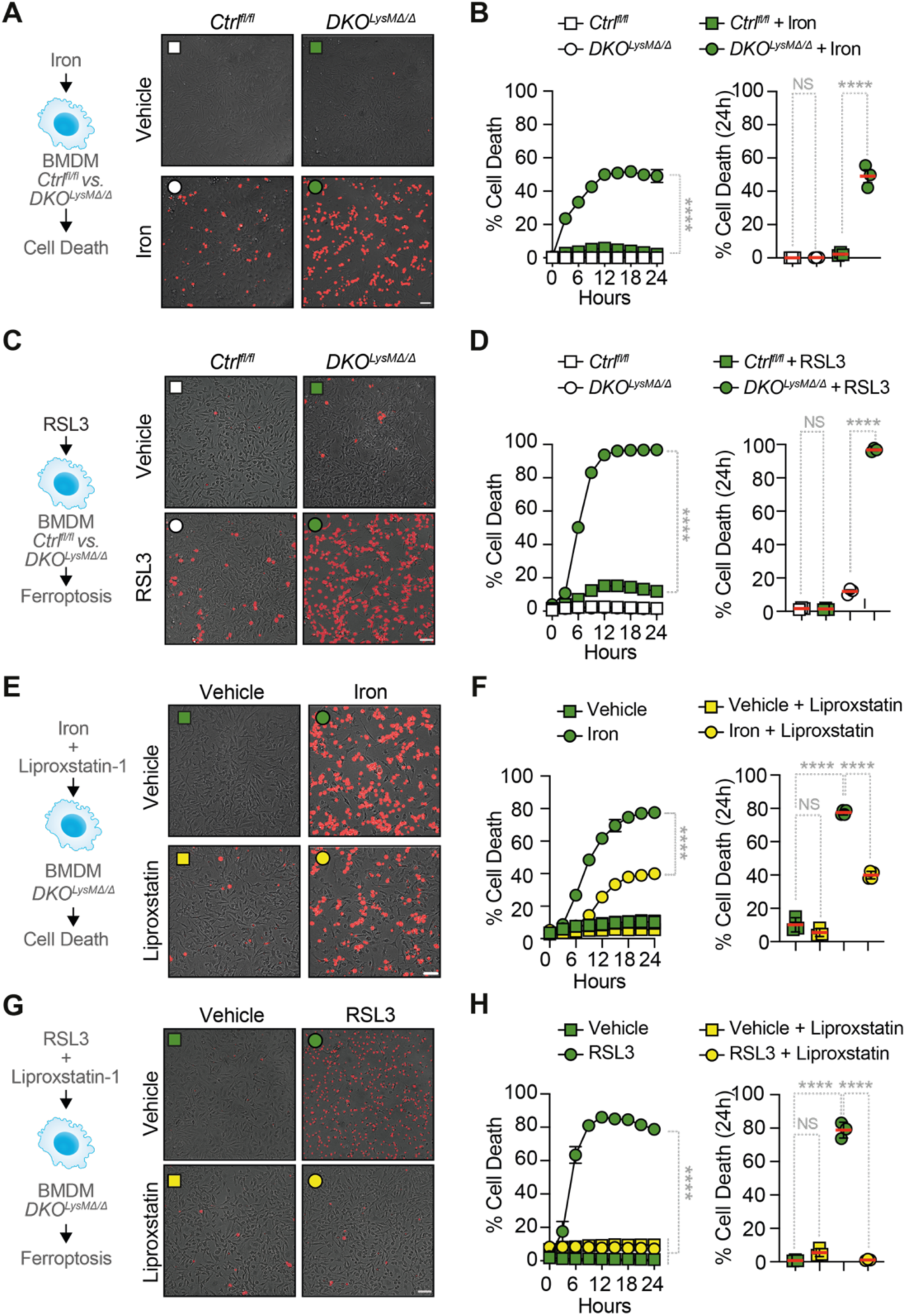
NRF2 and BVRA provide an integrated cytoprotective response against ferroptosis. (A and. **B)** Representative microscopy images (A) and quantification (B) of cell death (red, PI-positive cells) in *Ctrl^fl/fl^* and *DKO^LysMΔ/Δ^* BMDM treated with iron (ferric ammonium citrate; 200µM for 24h). (**C and D**) Representative microscopy images (**C**) and quantification (**D**) of cell death *Ctrl^fl/fl^* and *DKO^LysMΔ/Δ^* BMDM treated with RSL3 (125nM for 24h). (**E and F**) Representative microscopy images (**E**) and quantification **(F)** of cell death in *DKO^LysMΔ/Δ^* BMDM treated with iron in the presence or absence of liproxstatin-1 (Lip-1; 200nM for 24h). (**G and H**) Representative microscopy images (**G**) and quantification (**H**) of cell death in *DKO^LysMΔ/Δ^* BMDM treated with RSL3 in the presence or absence of liproxstatin-1. Data in (B), (D), (F), and (H) are presented as individual data points with bar graphs showing mean ± SEM; n=3 technical replicates, representative of at least two independent experiments. Scale bars in (A), (C), (E), and (G) = 50 μm. NS, not significant; ****p < 0.0001 by two-way ANOVA with Tukey’s multiple comparisons test.

To test whether the NRF2-independent axis that protects macrophages from ferroptosis relies on bilirubin production by BVRA^1^, we generated *Blvra^LysMΔ/Δ^* mice, carrying a *Blvra* deletion specifically in myeloid cells, as confirmed in BMDM by RT-qPCR and western blot (*Fig S4A-B*). Iron overload (*Fig S4C,D*) or inhibition of GPX4 by RSL3 (*Fig S4E,F*) failed to induce ferroptosis in *Blvra^LysMΔ/Δ^*BMDM, similar to control *Blvra^fl/fl^* BMDM (*Fig S4C-F*).

We hypothesized that NRF2 and BVRA are each sufficient per se to protect BMDM from ferroptosis. To test this hypothesis, we generated *Nrf2/Blvra^LysMΔ/Δ^*mice carrying a deletion of both *Nrf2* and *Blvra* in the myeloid compartment. Gene deletion in BMDM from *Nrf2/Blvra^LysMΔ/Δ^* mice, referred hereafter as double knockouts (*DKO^LysMΔ/Δ^*), was confirmed by RT-qPCR and western blot (*Fig. S5A-E*).

Iron overload (*Fig. 2A,B*) or RSL3 (*Fig. 2C,D*) triggered ferroptosis in *DKO^LysMΔ/Δ^*BMDM, under sub-cytotoxic conditions to control *Nrf2^fl/fl^Blvra^fl/fl^*(*Ctrl^fl/fl^*) BMDM (*Fig. 2A-D*). Liproxstatin-1 (Lip-1), a pharmacologic lipophilic RTA that suppresses ferroptosis^5^, protected *DKO^LysMΔ/Δ^* BMDM from ferroptosis, induced by iron overload (*Fig. 2E, F*) or RSL3 (*Fig. 2G,H*). Moreover, iron chelation by deferiprone protected *DKO^LysMΔ/Δ^* BMDM from ferroptosis, induced by RSL3 (*Fig. S5F,G*). This reveals that NRF2 and BVRA act in a coordinated and redundant manner to protect macrophages from ferroptosis.

### The cytoprotective effect of NRF2 and BVRA is specific to ferroptosis (*Fig.S6,7,8*)

We asked whether the cytoprotective effect of NRF2 and BVRA is exerted against other programmed cell death pathways, namely, apoptosis, necroptosis and pyroptosis. Pharmacologic inhibition of caspase 3, 7, and 8 by z-VAD-FMK^27^ failed to protect *DKO^LysMΔ/Δ^* BMDM from RSL3-induced ferroptosis (*Fig. S6A,B*). Induction of apoptosis across a range of Staurosporine concentrations^8^ was similar in *DKO^LysMΔ/Δ^ vs. Ctrl^fl/fl^*BMDM (*Fig. S6C,D*). This suggests that NRF2 and BVRA are not cytoprotective against caspase-dependent programmed cell death via apoptosis.

Pharmacologic inhibition of necroptosis by Necrostatin-1 (Nec-1), a selective inhibitor of the receptor-interacting protein kinase 1 (RIPK1)^11–13^, failed to protect *DKO^LysMΔ/Δ^* BMDM from ferroptosis (*Fig. S7A,B*). Induction of necroptosis by tumor necrosis factor (TNF) plus Second Mitochondria-Derived Activator of Caspases (SMAC) mimetics and z-VAD-FMK^28^ was also similar in *DKO^LysMΔ/Δ^ vs. Ctrl^fl/fl^* BMDM (*Fig. S7C,D*). This suggests that NRF2 and BVRA are not cytoprotective against RIPK1-dependent necroptosis.

Induction of programmed cell death by pyroptosis^9,10^, using a combination of bacterial lipopolysaccharide (LPS) and nigericin^9^, was also similar in *DKO^LysMΔ/Δ^ vs. Ctrl^fl/fl^* BMDM (*Fig. S8A,B*). This suggests that NRF2 and BVRA are not cytoprotective against pyroptosis.

Taken together, these data support the notion that NRF2 and BVRA are part of a coordinated cytoprotective program that protects macrophages specifically from ferroptosis, without interfering with other programmed cell death pathways.

### Glutathione and bilirubin mediate NRF2 and BVRA cytoprotection **(*Fig. 3*)**

Iron overload (*Fig.3A*) or inhibition of GPX4 by RSL3 (*Fig. 3B*) induced ROS accumulation in *DKO^LysMΔ/Δ^* BMDM, at concentrations that failed to induce ROS accumulation in control *Ctrl^fl/fl^* BMDM (*Fig. 3A,B*). Exogenous glutathione protected *DKO^LysMΔ/Δ^* BMDM from ferroptosis, induced by RSL3 (*Fig. 3C, D*), suggesting that glutathione provides a redundant cytoprotective effect against ferroptosis in the absence of bilirubin production by BVRA.

**Figure 3.**
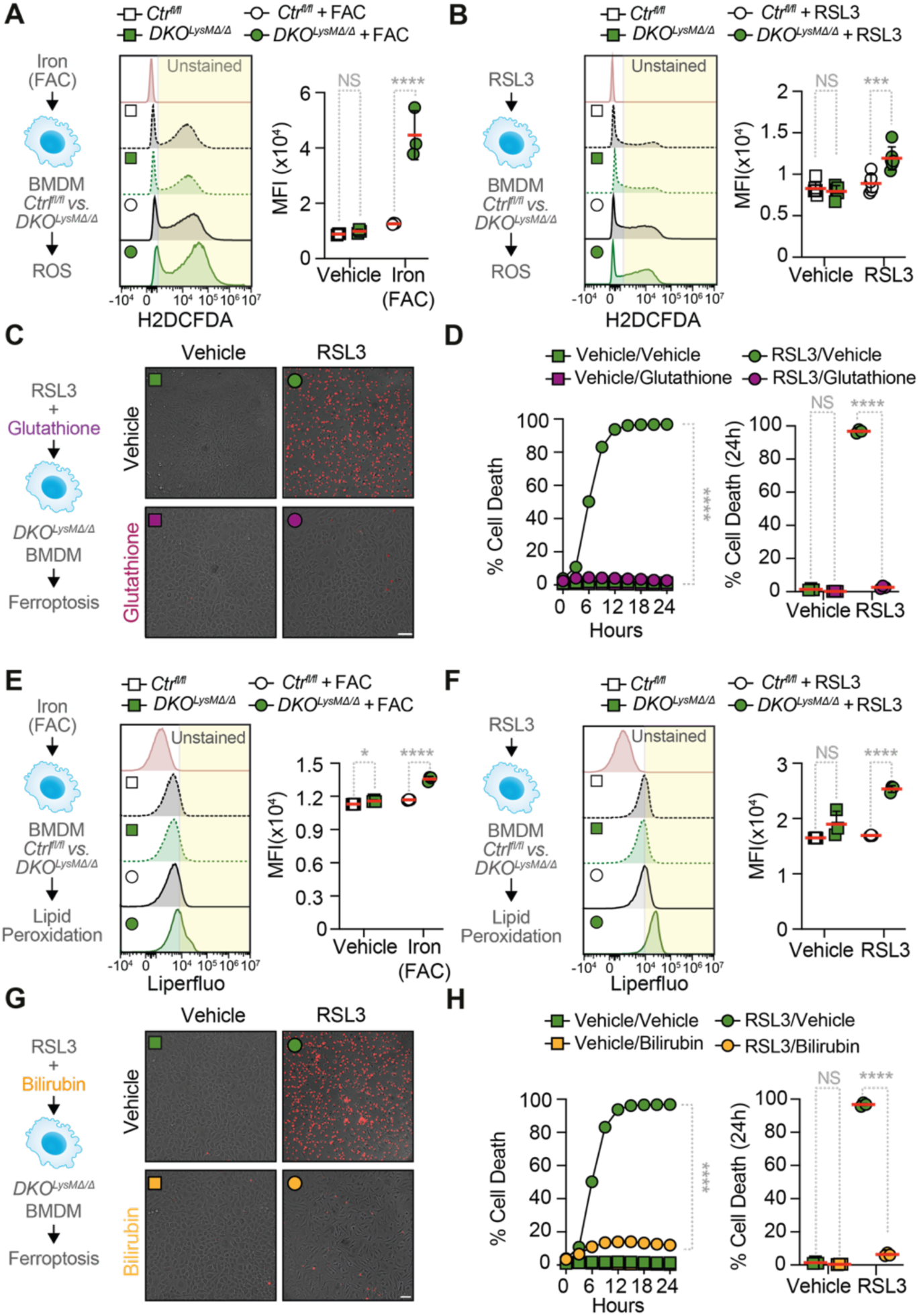
Glutathione and bilirubin are the downstream effectors of NRF2 and BVRA cytoprotection. (A and. **B)** Representative flow cytometry histograms and quantification of intracellular ROS in *Ctrl^fl/fl^* and *DKO^LysMΔ/Δ^* BMDM treated with (**A**) iron (ferric ammonium citrate; 200 µM for 6h) or **(B)** RSL3 (125 nM for 6h). Unstained controls are shown in pink. **(C and D)** Representative (**C**) microscopy images and (**D**) quantification of cell death (red, PI-positive cells) in *DKO^LysMΔ/Δ^* BMDM treated with RSL3 (125 nM for 24h) in the presence or absence of glutathione (5 mM for 24h). **(E and F)** Representative flow cytometry histograms and quantification of lipid peroxidation in *Ctrl^fl/fl^* and *DKO^LysMΔ/Δ^* BMDM treated with (**E**) iron (ferric ammonium citrate; 200 µM for 6h) or (**F**) RSL3 (125 nM for 6h). Unstained controls are shown in pink. **(G and H)** Representative (**G**) microscopy images and (**H**) quantification of cell death (red, PI-positive cells) in *DKO^LysMΔ/Δ^* BMDM treated with RSL3 (125 nM for 24h) in the presence or absence of bilirubin (200 nM for 24h). Data in (A), (B), (D), (E), (F) and (H) are presented as individual data points with bar graphs showing mean ± SEM. n=3 technical replicates representative of at least two independent experiments. Data in (C) and (G) show scale bars = 50 μm. NS, not significant; *p < 0.05, ***p < 0.001, ****p < 0.0001 by two-way ANOVA with Tukey’s multiple comparisons test.

Iron overload (*Fig.3E*) or GPX4 inhibition by RSL3 (*Fig. 3F*) induced lipid peroxidation in *DKO^LysMΔ/Δ^* BMDM, at concentrations that failed to induce lipid peroxidation in control *Ctrl^fl/fl^* BMDM (*Fig. 3E,F*). Exogenous bilirubin protected *DKO^LysMΔ/Δ^* BMDM from undergoing ferroptosis in response to RSL3 (*Fig. 3G,H*). This suggests that bilirubin provides a redundant cytoprotective effect against ferroptosis under glutathione depletion.

These observations suggest that impairing glutathione synthesis does not compromise cytoprotection as long as bilirubin remains available, and impaired bilirubin production also does not compromise cytoprotection as long as glutathione is available. Concurrent loss of both arms is therefore required to sensitize macrophages to ferroptosis.

### NRF2 and BVRA are required to sustain *in vivo* erythrophagocytosis (*Fig. 4, S9*)

To test whether NRF2 and BVRA are required to support erythrophagocytosis, we established an *in vivo* erythrophagocytosis assay in C57BL/6J mice receiving PKH26-labeled RBCs (*i.p.*). Within 3 hours, ∼50% of large peritoneal macrophages (LPM) phagocytosed RBC (PKH26^+^), reaching ∼80% at 17 and 24 hours (*Fig. 4A-C*). In contrast, small peritoneal macrophages (SPM) exhibited lower erythrophagocytosis, with ∼5%, 12%, and 17% PKH26^+^ cells at respective timepoints, maintaining near-baseline absolute numbers throughout (*Fig. 4A-C, S9A*). Ly6C^hi^ monocytes also showed minimal erythrophagocytic capacity, with 0.3%, 3%, and 2% PKH26^+^ cells and absolute numbers remaining at baseline across all timepoints (*Fig. 4A-C, S9A*). This shows that LPM are the dominant peritoneal erythrophagocytic macrophage population^29^.

**Figure 4.**
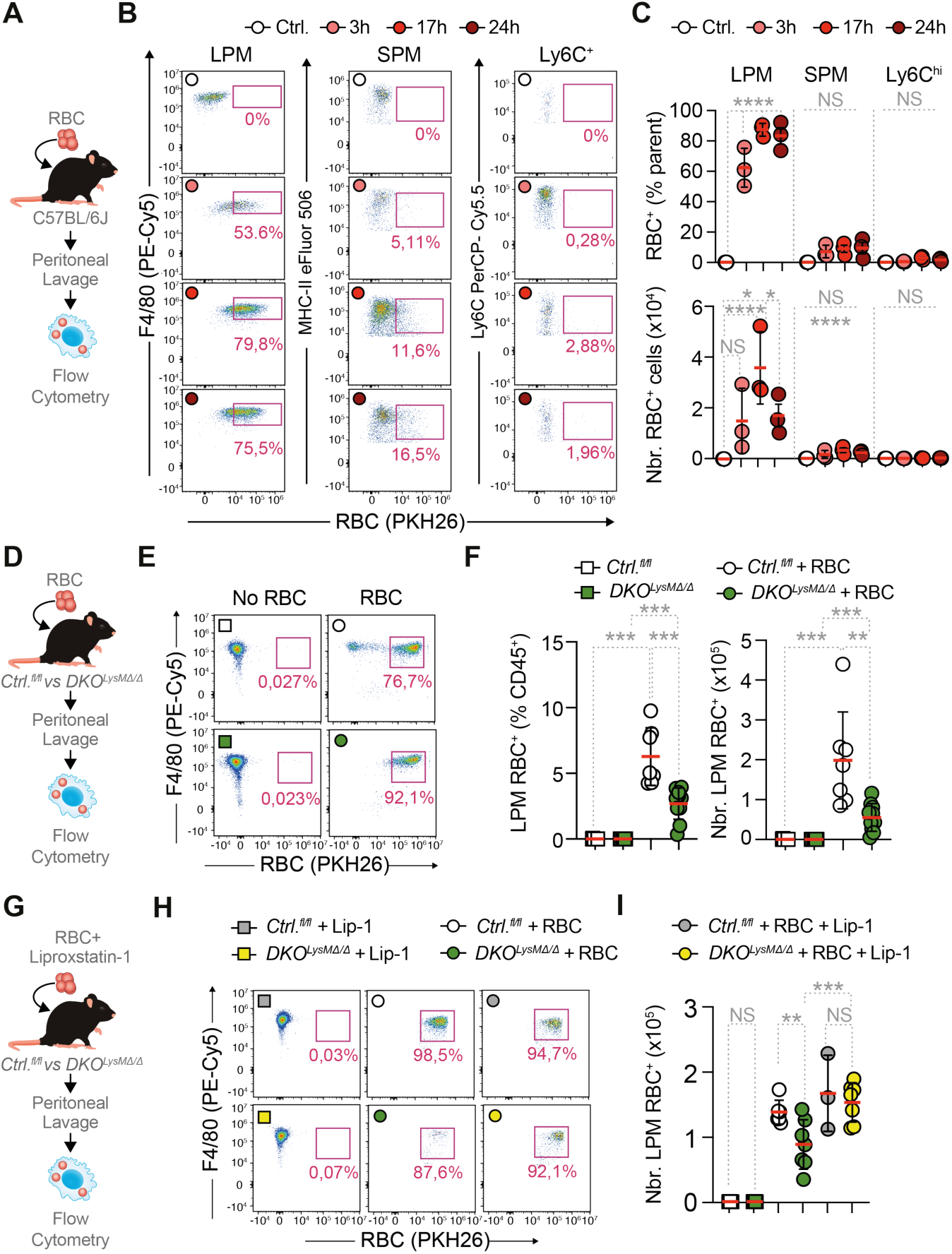
**NRF2 and BVRA decouple erythrophagocytosis from ferroptosis**. (**A**) Schematic representation of flow cytometric analysis of peritoneal myeloid populations in C57BL/6J mice receiving PKH26-labeled RBCs (*i.p*.). (**B and C**) Representative flow cytometry dot plots (**B**) and quantification of PKH26⁺ (RBC uptake) cells among large peritoneal macrophages (LPM; TIM-4⁺F4/80^hi^), small peritoneal macrophages (SPM; MHC-II⁺ F4/80⁺), and Ly6C⁺ monocytes at different time points post-injection. (**C**) Frequency (% of parent population, upper panel) and absolute counts (lower panel) of PKH26⁺ cells within each gated population are shown. (**D**) Schematic representation of flow cytometric analysis of LPM in *Ctrl^fl/fl^* and *DKO^LysMΔ/Δ^* mice receiving PKH26-labeled RBCs (i.p.). (**E and F**) Representative flow cytometry dot plots (**E**) and quantification of PKH26⁺ LPMs in *Ctrl^fl/fl^* and *DKO^LysMΔ/Δ^* mice before and after RBC injection (*i.p.*) (**F**). Frequency (% CD45⁺) and total peritoneal LPM numbers of erythrophagocytic LPMs. (**G**) Schematic representation of flow cytometric analysis of LPM in *Ctrl^fl/fl^* and *DKO^LysMΔ/Δ^* mice receiving PKH26-labeled RBCs (*i.p.*) and treated with liproxstatin-1 *vs.* vehicle. (**H and I**) Representative flow cytometry dot plots (**H**) and quantification of PKH26⁺ LPMs before and after RBC injection (*i.p.*) (**I**). Data in (C) and (F) are presented as individual data points showing mean ± SD. Data in (C) are from one experiment, n=3 mice *per* condition. Data in (F) and (I) are pooled from three independent experiments, n=7-10 mice *per* condition. NS, not significant; *p < 0.05, **p < 0.01, ***p < 0.001, ****p < 0.0001 by one-way or two-way ANOVA with Tukey’s multiple comparisons test.

Upon challenge with PKH26-labeled RBCs (*i.p*.), ∼92 and ∼77% of LPM contained intracellular RBC (PKH26^+^) within 24 hours, in *DKO^LysMΔ/Δ^* and *Ctrl^fl/fl^* mice, respectively (*Fig. 4D-F, S9B,C*). However, *DKO^LysMΔ/Δ^* mice presented a decrease greater than 50% in the total number of PKH26^+^ LPM compared to *Ctrl^fl/fl^* mice (∼2×10⁵ *vs.* ∼0.5×10⁵ cells, respectively) (*Fig. 4D-F, S9B,C*). This was accompanied by an increase in Ly6C^hi^ monocytes likely to repopulate the empty LPM niche (*Fig. S9B,C*). This suggests that NRF2 and BVRA are essential to support erythrophagocytosis, presumably protecting macrophages from undergoing ferroptosis.

### NRF2 and BVRA decouple erythrophagocytosis from ferroptosis **(*Fig. 4*)**

To determine whether the reduction in the number of PKH26^+^ LPM observed in *DKO^LysMΔ/Δ^*mice is attributable to ferroptosis, we performed the *in vivo* erythrophagocytosis assays under pharmacologic inhibition of ferroptosis by Lip-1. The reduction of PKH26^+^ LPM in *DKO^LysMΔ/Δ^ vs. Ctrl^fl/fl^* mice (*Fig. 4G-I*) was prevented by Lip-1 administration, rescuing the ∼50% deficit observed in the absence of ferroptosis inhibition (*Fig. 4G-I*). Lip-1 had no significant effect on PKH26^+^ LPM numbers in *Ctrl^fl/fl^*mice (*Fig. 4G-I*). This shows that the reduction of erythrophagocytic LPM in *DKO^LysMΔ/Δ^* mice is attributed functionally to ferroptosis and establishes NRF2 and BVRA as redundant cytoprotective mechanisms required for macrophages to withstand the oxidative burden imposed by erythrophagocytosis.

### NRF2 and BVRA define a ferroptosis-suppression program in RPM **(*Fig. 5, S10*)**

To determine whether NRF2 and BVRA operate in RPM we profiled splenic CD45⁺CD64⁺ mononuclear phagocytes from *Ctrl^fl/fl^* and *DKO^LysMΔ/Δ^* mice by single-cell RNA sequencing (scRNAseq) (*Fig. 5A, Fig. S10A*). Unsupervised clustering resolved the expected myeloid compartment, including classical and patrolling monocytes, dendritic cells, NF-ΚB-activated and stress-response macrophages, pre-RPM monocyte-derived macrophages, and a continuum of erythrophagocytic populations spanning early, mid, and late transitional RPMs through to mature RPM, based on *Spic* expression (*Fig. 5A,B; Fig. S10B*). Cluster composition was broadly preserved between genotypes (*Fig. S10C*). *Spic* expression rose progressively across the erythrophagocytic lineage, reaching its highest levels in mature RPMs (*Fig. S10D-F*). The RPM maturation-associated gene program was resolved at single-cluster resolution (*Fig. S10G*).

**Figure 5.**
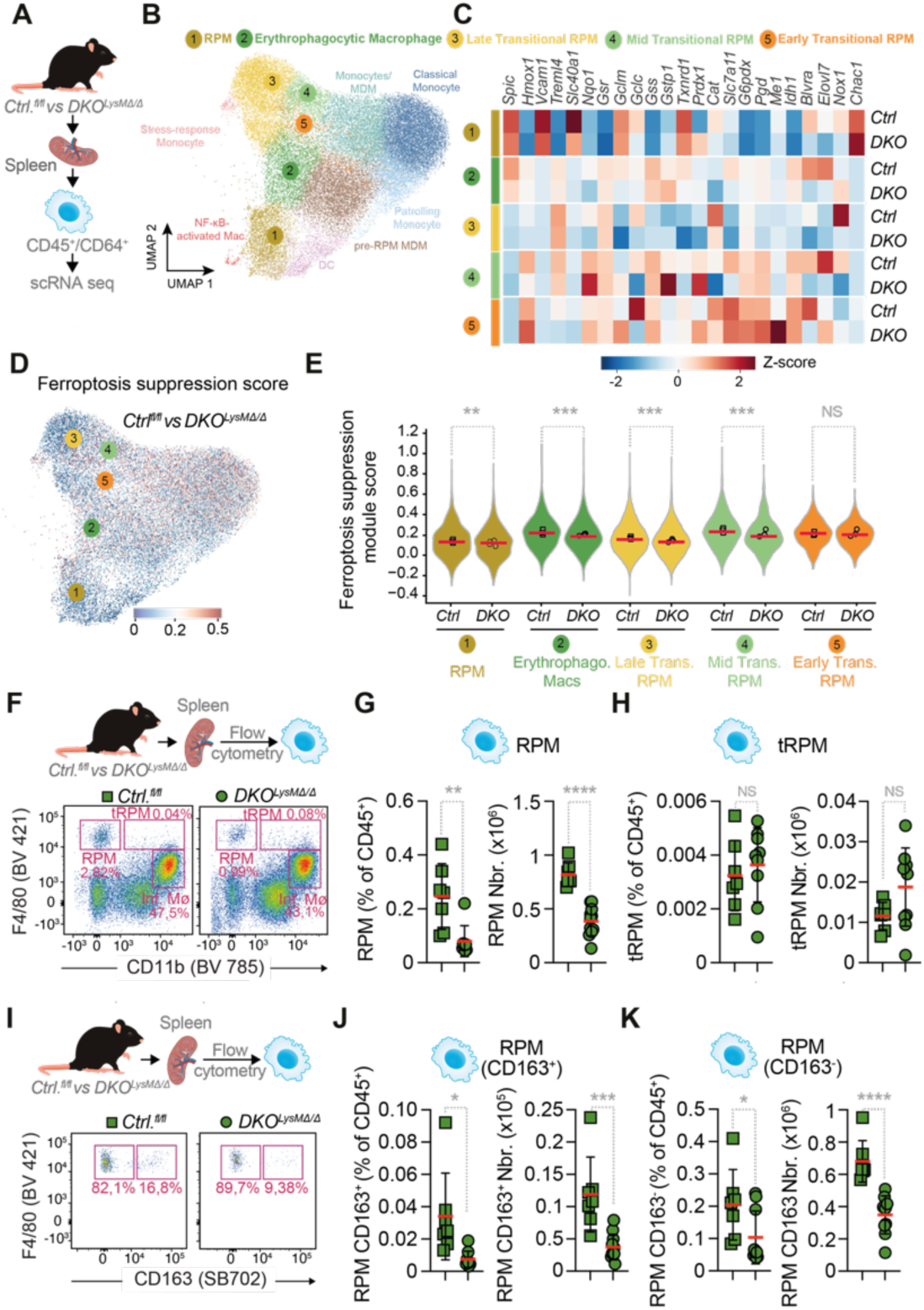
NRF2 and BVRA define a maturation-linked ferroptosis-suppression program in erythrophagocytic macrophages. (**A**) Splenic CD45⁺CD64⁺ cells were sorted from *Ctrl^fl/fl^*and *DKO^LysMΔ/Δ^* mice and subjected to single-cell RNA sequencing (scRNA-seq). (**B**) UMAP plot of scRNA-seq clusters from *Ctrl^fl/fl^*and *DKO^LysMΔ/Δ^* spleens, colored by cell population identity (total n=37,739 22,221 cells). (**C**) Heatmap of selected marker gene expression (Z-score) across macrophage clusters and genotypes from scRNA-seq. (**D**) UMAP projection of ferroptosis suppression module score in *Ctrl^fl/fl^* and *DKO^LysMΔ/Δ^* splenic macrophages defined *a priori* from FerrDb v2^30^ and KEGG pathway mmu04216. (**E**) Violin plots of ferroptosis suppression module score across macrophage subpopulations in *Ctrl^fl/fl^* and *DKO^LysMΔ/Δ^* mice. (**F**) Representative flow cytometry dot plots of splenic myeloid populations from *Ctrl^fl/fl^* and *DKO^LysMΔ/Δ^* mice, gated on live CD45⁺Ly6G⁻CD11c⁻ cells and identified by F4/80 and CD11b expression. Red pulp macrophages (RPM), transitional RPM (tRPM), and inflammatory monocytes/macrophages (Inf. Mø) are indicated. (**G**) Frequency (% of CD45⁺) and absolute numbers of RPM in *Ctrl^fl/fl^* and *DKO^LysMΔ/Δ^* spleens. (**H**) Frequency (% of CD45⁺) and absolute numbers of tRPM in *Ctrl^fl/fl^* and *DKO^LysMΔ/Δ^* spleens. (**I**) Representative flow cytometry dot plots of CD163 expression among RPMs (F4/80 *vs*. CD163) in *Ctrl^fl/fl^* and *DKO^LysMΔ/Δ^* spleens. (**J**) Frequency (% of CD45⁺) and absolute numbers of CD163⁺ RPM in *Ctrl^fl/fl^*and *DKO^LysMΔ/Δ^* spleens. (**K**) Frequency (% of CD45⁺) and absolute numbers of CD163⁻ RPM in *Ctrl^fl/fl^* and *DKO^LysMΔ/Δ^* spleens. Data in (B-E), n=3-4 animals *per* condition. Data in (G), (H), (J), and (K) are presented as individual data points showing mean ± SD, pooled from at least three independent experiments, n=7-11 mice *per* condition. Statistical comparisons in (E) were performed using a Wilcoxon rank-sum test. NS, not significant; *p < 0.05, **p < 0.01, ***p < 0.001, ****p < 0.0001 by two-tailed unpaired t-test.

The RPM core program (*Spic*, *Hmox1*, *Blvra*, *Vcam1*, *Treml4*, *Slc40a1*), NRF2 antioxidant network (*Nqo1*, *Gsr*, *Gclm*, *Gclc*, *Gss*, *Gstp1*, *Txnrd1*, *Prdx1*, *Cat*, *Slc7a11*) and NADPH-generating enzymes (*G6pdx*, *Pgd*, *Me1*) were all selectively downregulated in *DKO^LysMΔ/Δ^ vs. Ctrl^fl/fl^*mature RPMs (cluster 1) and to a graded extent across the transitional populations (clusters 3, 4). In contrast, monocyte-derived erythrophagocytic macrophages (cluster 2) showed comparatively modest changes (*Fig.5C*).

A composite ferroptosis-suppression score defined from FerrDb v2^30^ and KEGG pathway mmu04216 mapped most strongly onto the RPM and transitional RPM clusters and was significantly reduced in *DKO^LysMΔ/Δ^ vs. Ctrl^fl/fl^*mice across all four clusters (1-4), with the earliest transitional population (cluster 5) preserved (*Fig. 5D,E*). The ferroptosis-suppression score was also significantly reduced in monocyte/MDM and pre-RPM monocyte-derived macrophage populations, but not in other myeloid subsets (*Fig. S10H*). The transcriptional defect thus tracks the maturation trajectory of erythrophagocytic macrophages, intensifying as these acquire the full RPM identity. These findings establish that the NRF2/BVRA ferroptosis-control program is not a generic myeloid stress response, but a lineage-specific adaptation selectively engaged through erythrophagocytic commitment.

### NRF2 and BVRA are required to maintain the RPM compartment (*Fig. 5, S11,12*)

To determine whether NRF2 and BVRA are required for the maintenance of RPM *in vivo*, splenic RPM (Lin⁻Ly6G⁻CD11c⁻F4/80^hi^CD11b^low^) were immunophenotyped by flow cytometry at 16 weeks of age. *DKO^LysMΔ/Δ^* mice presented roughly three-fold fewer mature splenic RPM than *Ctrl^fl/fl^* mice, whether expressed as a fraction of CD45⁺ cells or as absolute number (*Fig. 5F,G*). Transitional RPM (tRPM) were unaffected in both frequency and number (*Fig. 5H*), placing the cellular loss at the RPM maturation step rather than at earlier progenitors.

Dissection of the mature RPM compartment by CD163 expression^10^ revealed that both the CD163⁺ subset, associated with extracellular Hb handling, and the CD163⁻ subset was depleted in *DKO^LysMΔ/Δ^ vs. Ctrl^fl/fl^*mice at 16 weeks of age (*Fig. 5I-K*). This reveals that the ferroptosis-control program enforced by NRF2 and BVRA is required for the survival of the entire mature RPM population rather than a single functional subset.

Analysis of single *Nrf2^LysMΔ/Δ^* or *Blvra^LysMΔ/Δ^*mice at 16 weeks of age showed no differences in RPM, RPM CD163⁺, or RPM CD163⁻ populations compared to respective genetic controls (*Fig. S11A-L*). This is consistent with the NRF2-dependent program compensating for the BVRA-dependent one, and vice versa, similar to identified for BMDM (*Fig. 2, Fig. 3*).

In contrast to RPM, other splenic myeloid populations, including inflammatory macrophages, dendritic cells, and neutrophils, were not significantly altered in *DKO^LysMΔ/Δ^ vs. Ctrl^fl/fl^*mice at 16 weeks of age (*Fig. S12A-D*), suggesting a specific dependence of the mature RPM compartment on this antioxidant axis. Analysis of single *Nrf2^LysMΔ/Δ^* (*Fig. S12E-G*) or *Blvra^LysMΔ/Δ^* (*Fig. S12H-J*) mice also revealed no differences in other splenic myeloid populations, including inflammatory macrophages, dendritic cells, and neutrophils.

These observations establish the NRF2 and BVRA antioxidant pathways as critical axes to sustain the mature RPM compartment *in vivo*. Loss of both axes produces a graded transcriptional defect along the erythrophagocytic differentiation trajectory and a selective collapse of the terminally differentiated RPM compartment.

### NRF2 and BVRA act specifically in erythrophagocytic macrophages **(*Fig. S13*)**

The number of peritoneal LPM and SPM and their percentage among CD45⁺ cells were indistinguishable in *DKO^LysMΔ/Δ^ vs. Ctrl^fl/fl^* mice (*Fig. S13A,B*). The percentage of bone marrow macrophages (BMM) among CD45⁺ cells was increased without affecting their numbers in *DKO^LysMΔ/Δ^ vs. Ctrl^fl/fl^* mice (*Fig. S13C,D*). There was no change in the percentage or absolute numbers of Ly6C^hi^ monocytes in the bone marrow (*Fig. S13C,D*). The number of liver Kupffer cells (KC) was increased in *DKO^LysMΔ/Δ^ vs. Ctrl^fl/fl^* mice (*Fig. S13E,F*). Transient monocytes known to participate in stress erythrophagocytosis^23^ were likewise unaffected (*Fig. S13E,F*). Lung alveolar macrophage (AM) numbers were decreased in *DKO^LysMΔ/Δ^ vs. Ctrl^fl/fl^* mice (*Fig. S13G,H*) while the number of lung interstitial macrophages (IM) remained unaffected. Together, these data indicate that the NRF2/BVRA antioxidant axis is required for maintenance of the mature RPM lineage.

### Myeloid NRF2 and BVRA regulate organismal iron homeostasis and iron-deficiency anemia **(*Fig. 6*)**

To investigate whether the expression of NRF2 and BVRA in the myeloid compartment regulates systemic iron homeostasis we quantified splenic iron content under steady state conditions. *DKO^LysMΔ/Δ^*mice presented a marked accumulation of iron in the red pulp of the spleen, as compared to *Ctrl^fl/fl^* mice at 16 weeks of age (*Fig. 6A,B*). This is reminiscent of splenic iron accumulation in *Spic*-deficient mice consistent with RPM depletion^7,8^.

**Figure 6.**
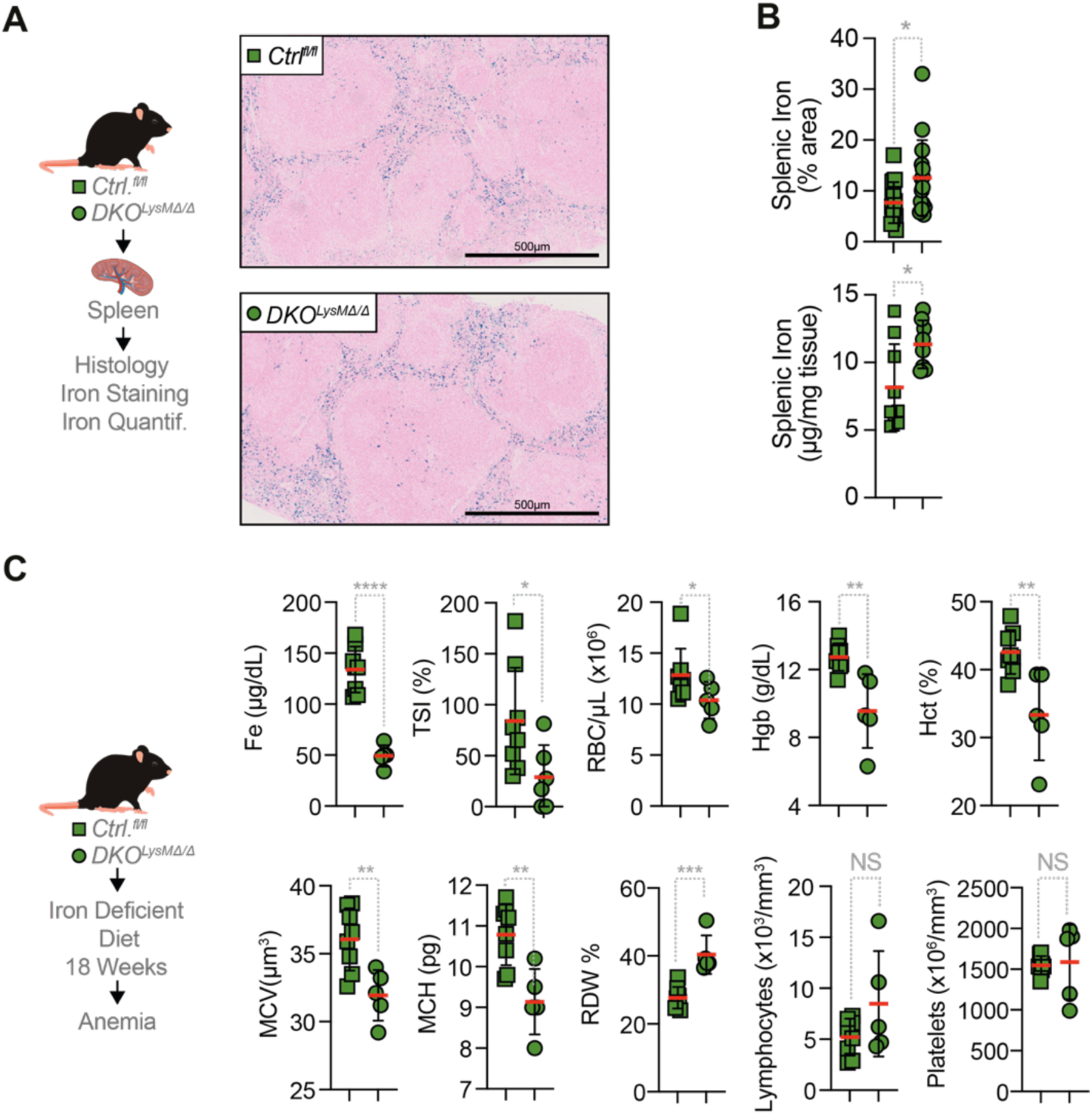
**NRF2 and BVRA expression by RPM regulates organismal iron homeostasis and prevents iron-deficiency anemia**. **(A)** Representative histological images of Prussian blue iron staining in spleens from *Ctrl^fl/fl^* and *DKO^LysMΔ/Δ^*mice. Scale bars as indicated. **(B)** Quantification of splenic iron as percentage of stained area (upper panel) and concentration *per* milligram of tissue (lower panel) in *Ctrl^fl/fl^*and *DKO^LysMΔ/Δ^* mice. **(C)** Experimental schematic and hematological parameters in *Ctrl^fl/fl^* and *DKO^LysMΔ/Δ^* mice fed an iron-deficient diet for 18 weeks: serum iron (Fe, µg/dL), red cell distribution width (RDW, %), red blood cell count (RBC/µL, ×10^6^), hemoglobin (Hgb, g/dL), hematocrit (Hct, %), mean corpuscular volume (MCV, µm^3^), mean corpuscular hemoglobin (MCH, pg), transferrin saturation index (TSI, %), Lymphocytes (×10^3^/mm^3^), platelets (×10^6^/mm^3^). Data in **(B)** are pooled from at least three independent experiments, n=8–16 mice *per* condition. Data in **(C)** are from two independent experiments, n=6-8 mice *per* condition. NS, not significant; *p < 0.05, **p < 0.01, ***p < 0.001, ****p < 0.0001 by two-tailed unpaired t-test or two-tailed Mann-Whitney test.

When maintained under an iron-deficient diet for 18 weeks *DKO^LysMΔ/Δ^*mice reduced serum iron concentration to 50-120 µg/dL, compared to 110-160 µg/dL in *Ctrl^fl/fl^* mice (*Fig. 6C*). Concomitantly, there was a reduction of transferrin saturation index (% of iron bound to transferrin) to 30% in *DKO^LysMΔ/Δ^* mice, compared to 85% in *Ctrl^fl/fl^* mice (*Fig. 6C*). This was not observed under standard (iron rich) diet (*Fig. S14A*), demonstrating that myeloid NRF2 and BVRA become essential to support systemic iron metabolism under dietary iron deficiency.

Under iron-deficient diet, *DKO^LysMΔ/Δ^* mice reduced circulating RBC count to 10,3×10^6^ µL, compared to 12,8×10^6^ µL in littermate *Ctrl^fl/fl^* mice (*Fig. 6C*). This was associated with a more pronounced decrease of Hb concentration and hematocrit (*Fig. 6C*). Mean corpuscular volume (MCV) and mean corpuscular hemoglobin (MCH) were also lower in *DKO^LysMΔ/Δ^ vs. Ctrl^fl/fl^* mice, consistent with microcytic iron-deficiency anemia. This was accompanied by increased RBC distribution width (RDW) in *DKO^LysMΔ/Δ^ vs. Ctrl^fl/fl^* mice indicating the development of anisocytosis, while lymphocyte and platelet numbers remained unaffected, demonstrating that this phenotype is selective to erythroid parameters (*Fig. 6C*). This was not observed under standard (iron rich) die (*Fig. S14A*), demonstrating that myeloid NRF2 and BVRA become essential to the severity of iron-deficient anemia, highlighting an essential macrophage-dependent pathway linking iron recycling and ferroptosis to RBC homeostasis.

## Discussion

Erythrophagocytic macrophages face a fundamental paradox whereby their effector function, which consists in extracting iron from Hb-bound heme in damaged RBC, generates a flux of intracellular catalytic iron that if not controlled can fuel lipid peroxidation and induce cell death via ferroptosis. We found that the heme-responsive transcription factor SPI-C resolves this paradox by coupling erythrophagocytosis to a robust and redundant anti-ferroptosis program. This transcriptional program is not required to support erythrophagocytic macrophage development from yolk sac progenitors, similar to described for *Spic*-deficient mice^7,8^. Instead, the anti-ferroptosis program regulated by SPI-C establishes a fail-safe mechanism for survival of erythrophagocytic macrophages, independently of their origin.

Consistent with SPI-C activation inducing a glutathione-dependent and-independent anti-ferroptosis axes (*Fig. 4G-I, Fig. S9*), genetic ablation of both axes renders macrophages susceptible to undergo ferroptosis, but only upon erythrophagocytosis (*Fig. 4G-I, Fig. S9*). This in turn depletes RPM (*Figs. 5F-K*) and compromises systemic iron recycling, dysregulating organismal iron homeostasis (*Figs. 4,5*) and exacerbating the severity of iron-deficiency anemia (*Figs. 6A-C*).

Upon activation by heme, SPI-C induces the expression of a number of NRF2-regulated genes, without however inducing *Nrf2* mRNA expression (*Fig. 1D-F*). A plausible mechanism might involve *Sqstm1* (encoding p62), one of the most strongly SPI-C-dependent heme-induced genes (*Fig. 1G*) recruited to erythrocyte-containing phagosomes^31^ to activate NRF2 via sequestration of its constitutive repressor KEAP1^32,33^. SPI-C activation also induces the transcription of NADPH-generating enzymes, which are required to support HO-1 and BVRA activity underlying bilirubin production (*Fig. 1G*). Bilirubin is a lipophilic RTA^34^ that protects macrophages from ferroptosis, backing up the glutathione-dependent GPX4 axis (*Fig. 3G-H*). This BVRA-bilirubin axis defines a distinct glutathione-independent regulator of ferroptosis, alongside other lipophilic RTA, such as ubiquinol generated by the NADPH-dependent oxidoreductases FSP1/AIFM2^24,25^. Unlike FSP1, BVRA is tailored to erythrophagocytosis, where heme catabolism generates both catalytic iron and biliverdin, scaling iron-mediated oxidative burden with ferroptosis protection. This metabolic self-regulation, in which a byproduct of heme catabolism fuels its own counterregulatory mechanism, represents a possible solution to overcome the oxidative demands of erythrophagocytosis and iron recycling by RPM.

NRF2 and BVRA do not interfere with other programmed cell death pathways, including apoptosis, necroptosis, or pyroptosis (*Fig. S6-8*). This is consistent with glutathione and bilirubin converging to limit PUFA peroxidation, the ferroptosis executioner, rather than upstream signaling transduction shared across other programmed cell death pathways.

The selective depletion of RPM, with increase of Kupffer cell and maintenance of bone marrow macrophages in *DKO^LysMΔ/Δ^* mice (*Fig. S13A-F*), establishes ferroptosis as a specific vulnerability of erythrophagocytic macrophages. Under steady state conditions RPM are uniquely exposed to a massive flux of intracellular catalytic iron, a major constraint not faced by other macrophage populations^35^.

A partial depletion of alveolar macrophages in *DKO^LysMΔ/Δ^*mice (*Fig. S13G,H*) is consistent with lipid peroxidation stress associated with lung surfactant clearance^36^. This is also consistent with the lipid sensor and nuclear transcriptional regulator Peroxisome Proliferator-Activated Receptor gamma (PPARγ) being critically involved in maintenance of both RPM and alveolar macrophages under steady state conditions^37^.

The systemic consequences of RPM ferroptosis in *DKO^LysMΔ/Δ^*mice phenocopy the age-dependent RPM depletion and splenic iron accumulation reflecting the loss of iron recycling in *Spic*-deficient mice^7,8^. *DKO^LysMΔ/Δ^* mice developed severe anemia under dietary iron restriction, where erythropoiesis becomes critically dependent on macrophage-mediated erythrophagocytosis and iron recycling^38^ (*Fig. 6C*). This establishes a causal link between ferroptosis and iron-deficiency anemia, whereby ferroptosis resistance enforces systemic iron homeostasis.

The clinical relevance of the anti-ferroptosis axis is consistent with the identification of Glutathione S-transferase Alpha 1 (*GSTA1*) among candidate iron-deficiency anemia susceptibility gene in a GWAS meta-analysis^39^. Multiple GSTA family members, including *Gsta3* and *Gsta4*, are among the most strongly heme-induced and SPI-C-dependent transcripts (*Fig. 1G*), providing human genetic support for the SPI-C-NRF2-glutathione axis in iron-deficiency anemia progression.

We propose that ferroptosis operates as a rate-limiting constraint on RPM with physiologic attrition normally offset by monocyte-derived replenishment. Under conditions of heightened erythrophagocytic demand, as demonstrated for ageing and iron deficiency, sustained intracellular catalytic iron flux induces RPM ferroptosis, impairing iron recycling and promoting anemia severity. Whether this also occurs under hemolytic conditions, including malaria, is likely but remains to be established.

In malaria, where Mendelian randomization establishes a causal link between infection and anemia attributed to hepcidin-mediated suppression of iron absorption^40^, our findings suggest a complementary mechanism. Namely, hepcidin-induced ferroportin degradation on RPM, traps intracellular iron amplifying ferroptotic pressure and limiting iron recycling, which promotes severe anemia. Presumably this explains why iron recycling by renal proximal tubule epithelial cells becomes rate limiting during malaria^41^. These considerations suggest that pharmacological strategies that enhance RPM ferroptosis resistance, through ferroptosis inhibitors or enhancement of the BVRA-bilirubin axis, could represent a mechanistically distinct therapeutic approach to iron deficiency anemia that targets the recycling machinery rather than iron supply.

In conclusion, SPI-C embeds ferroptosis resistance into erythrophagocytic macrophage function through coordinated induction of the NRF2-glutathione and BVRA-bilirubin axes. This dual protection system is essential for erythrophagocytic macrophage function, supporting iron recycling and systemic iron homeostasis. These findings identify bilirubin as a physiological ferroptosis inhibitor and nominate the SPI-C-NRF2-BVRA pathway as a potential therapeutic target for iron-deficiency anemia affecting approximately one-third of the global population.

### Limitations of the study

This study employs the LysM-Cre system, which achieves efficient gene deletion in a fraction of but not all myeloid cells, including macrophages. The study was performed in mice, and whether the SPI-C-NRF2-BVRA axis is operational in human RPM remains to be established. The relative contributions of bilirubin’s radical-trapping, mitochondrial, and Fenton-attenuating activities to ferroptosis protection, and its intracellular concentrations during active erythrophagocytosis, warrant further investigation.

## RESOURCE AVAILABILITY

### Lead contact

Requests for further information and resources should be directed to and will be fulfilled by the lead contact, Miguel P. Soares.

### Materials availability

Mouse strains generated in this study are available from the lead contact with a completed materials transfer agreement.

### Data and code availability

- All data reported in this paper will be shared by the lead contact upon request.
- Code developed in this paper will be shared by the lead contact upon request.
- Any additional information required to reanalyze the data reported in this paper is available from the lead contact upon request.

## Acknowledgments

The authors are indebted to all members of the Inflammation group (GIMM) for insightful technical and intellectual contributions, to the staff at the GIMM Histopathology (Pedro Faisca), Flow Cytometry, Rodent, Bioimaging, Genomics, Advanced Data Analysis (Hugo Lainé) and Metabolomics facilities.

## Author Contributions

Conceptualization: MPS

Formal analysis: MM and EJ

Resources: BDP

Investigation: MM, EJ, MP, MC, ST, RM, JZK, SC, MT, SV, AF, NBS, EM.

Visualization: MM and MPS

Funding acquisition: MPS

Project administration: MPS

Supervision: MPS and EJ

Writing: original draft: MPS

Writing: review & editing: MM

## Declaration of generative AI and AI-assisted technologies in the manuscript preparation process

During the preparation of this work the author(s) used AI-assisted technologies in the manuscript preparation process. Author(s) reviewed and edited the content as needed and take full responsibility for the content of the published article.

## Methods

### KEY RESOURCES TABLE

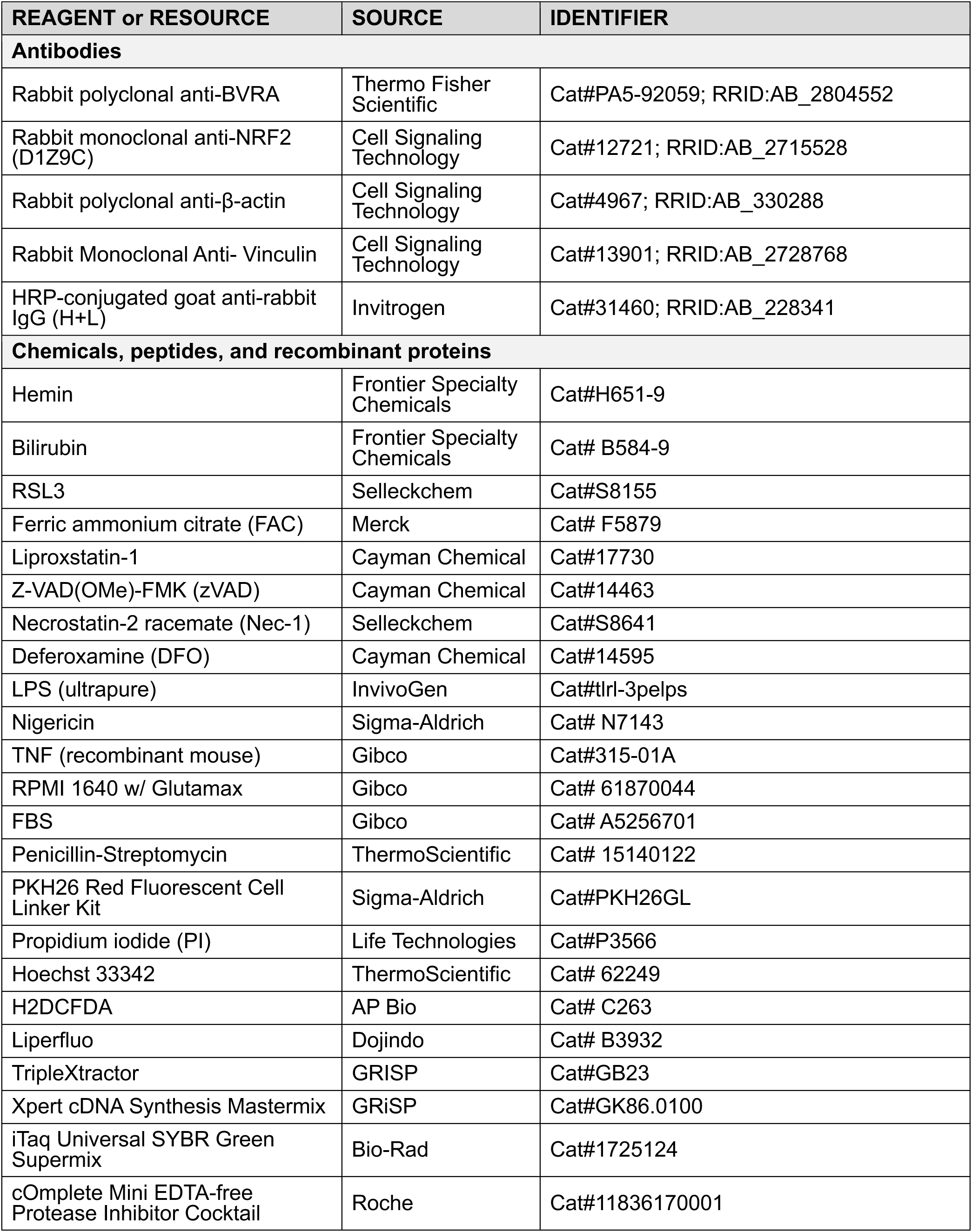
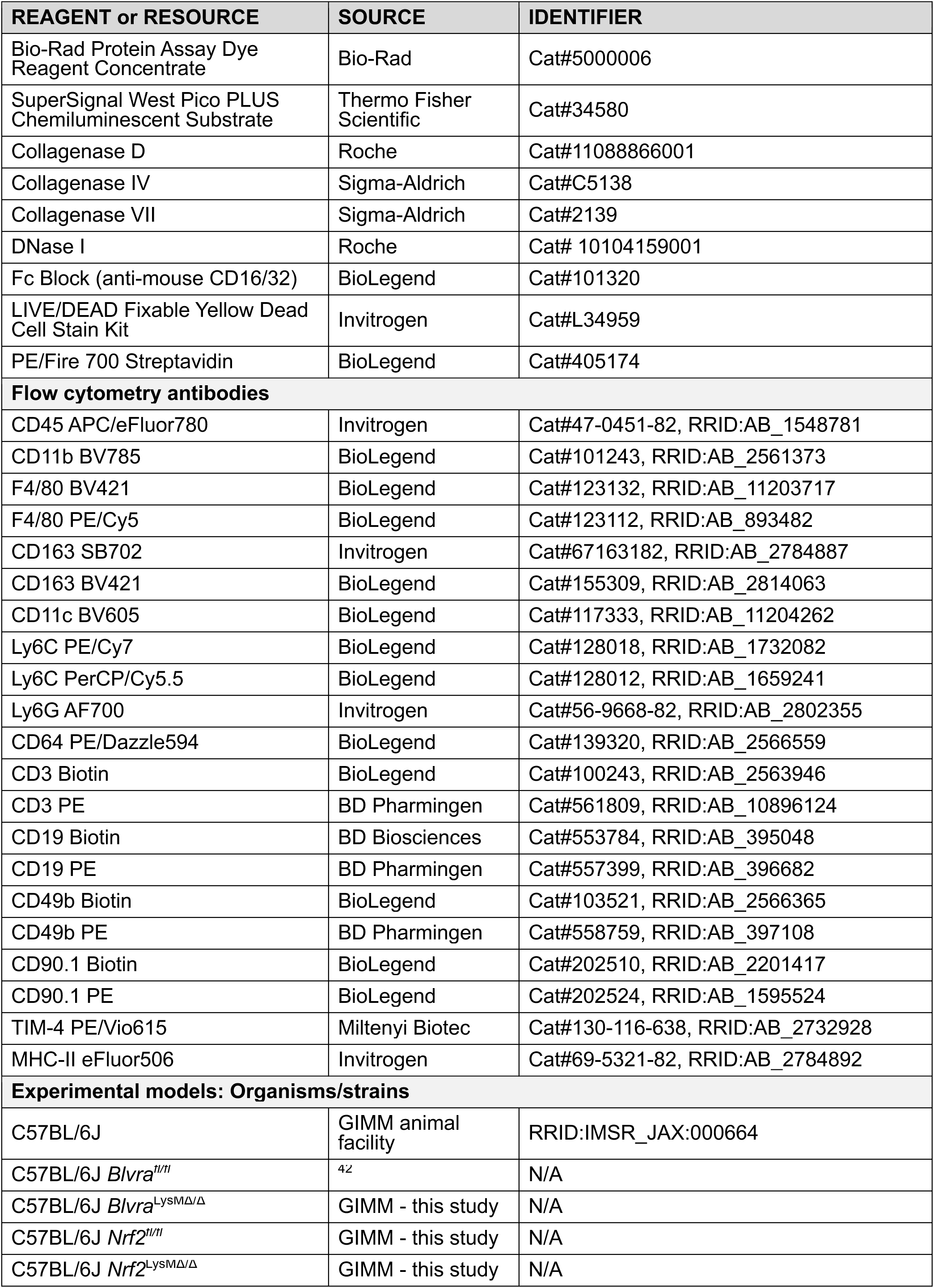
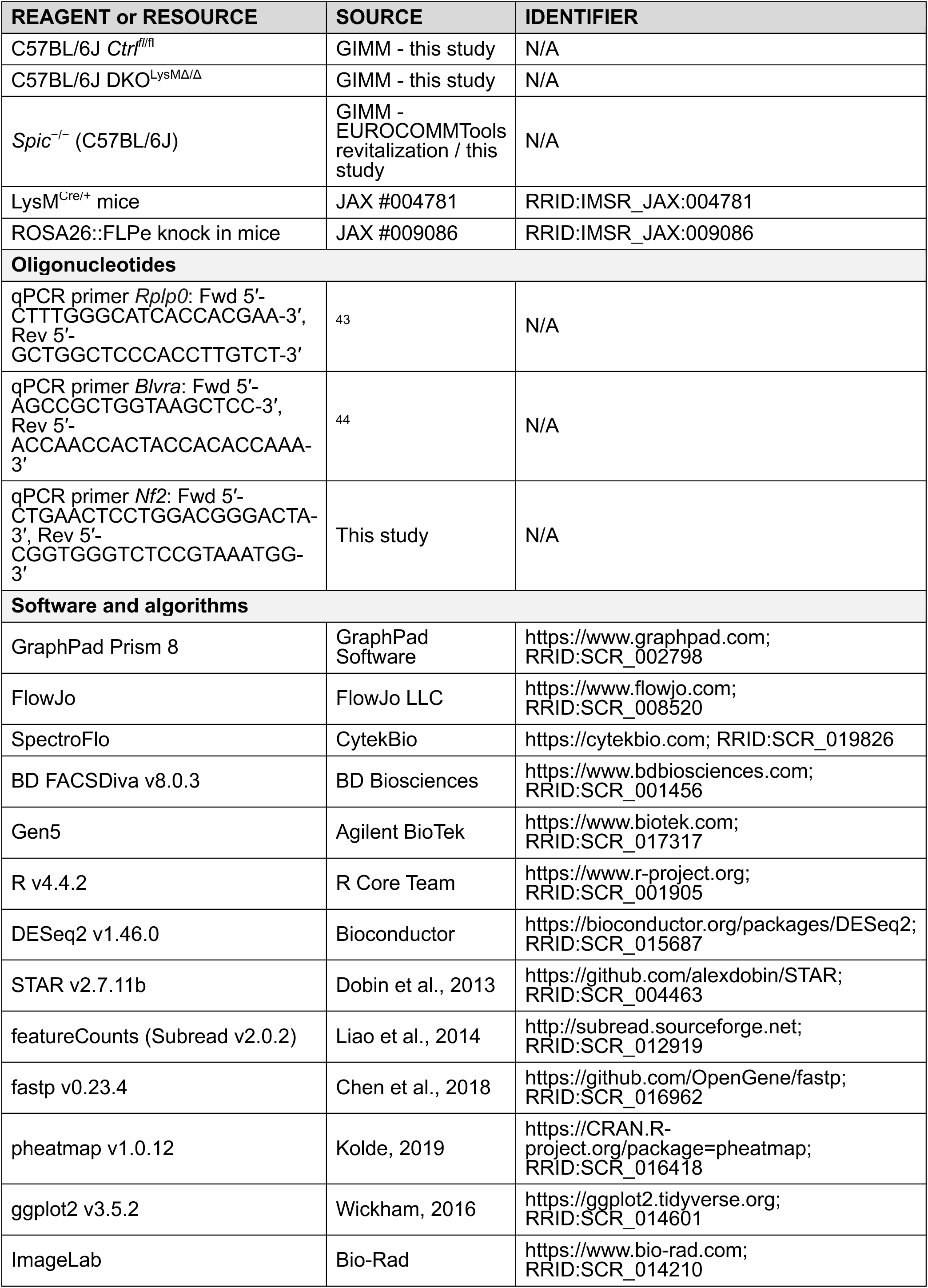
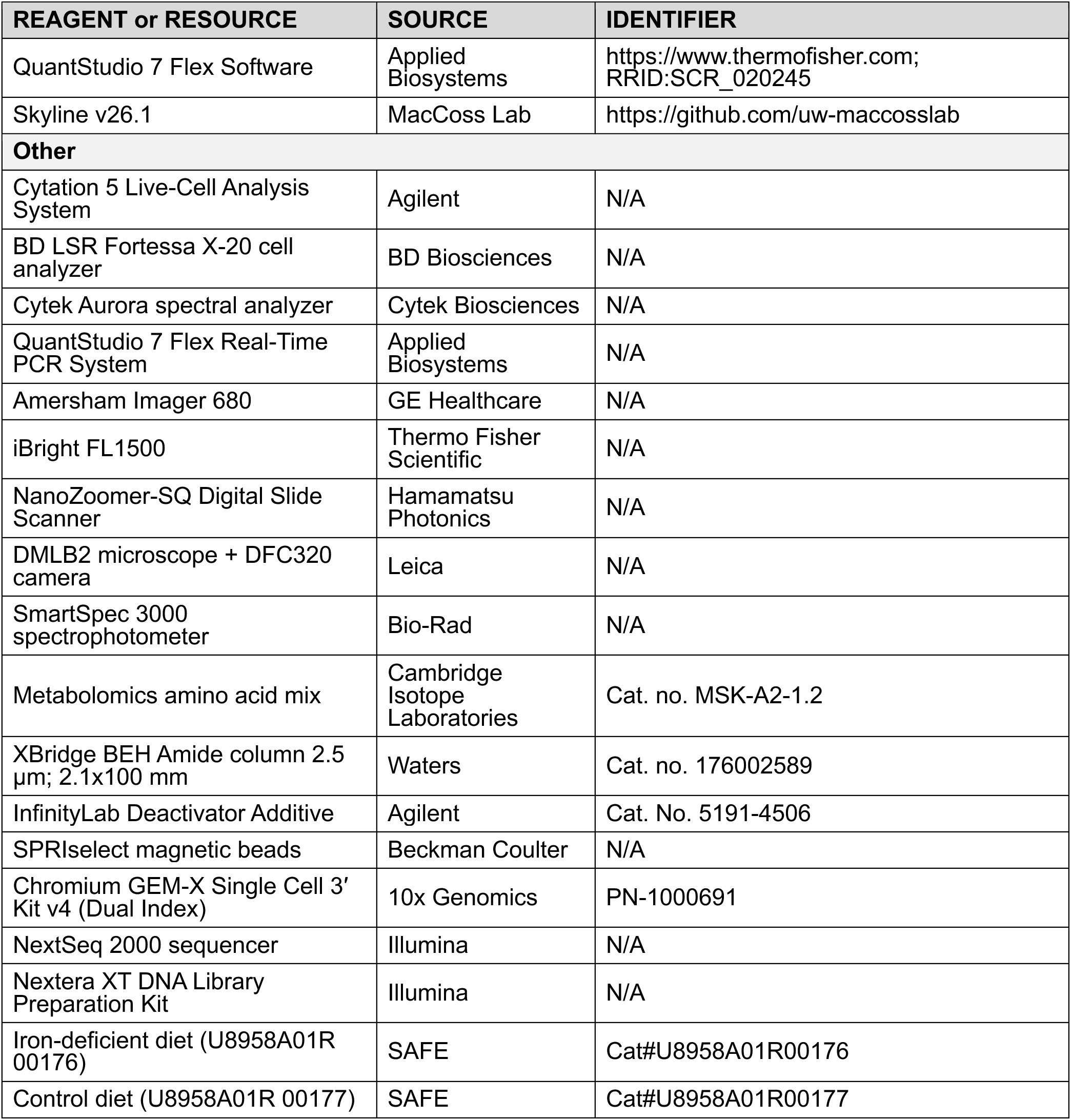

### Mice

Mice (male and female) were bred and maintained under specific pathogen-free (SPF) conditions at the Gulbenkian Institute for Molecular Medicine (GIMM). Mice were housed at standard *vivarium* temperature (22°C) in a 12-hour light/dark cycle with free access to water and standard chow pellets. All experimental protocols were approved in a two-step procedure, by the Animal Welfare Body of the GIMM and by the Portuguese National Entity that regulates the use of laboratory animals in research (Direção Geral de Alimentação e Veterinária; DGAV). Experimental procedures followed the Portuguese (Decreto-Lei no 113/2013) and European (Directive 2010/63/EU) legislation. All experiments were conducted between 12-16 weeks of age. C57BL/6J mice were obtained from GIMM animal facility. All animals were used in the C57BL/6J background. *Nrf2^fl/fl^*mice were generated at GIMM (Figure S3) by microinjecting the *Nfe2l2^tm1a(EUCOMM)Hmgu^* ES cell line (obtained from EUCOMM) into C57BL/6J blastocysts, which were then transferred into pseudo pregnant females. Male chimeras were screened for germline transmission by backcrossing to C57BL/6J females. F1 offspring heterozygous for the *Nfe2l2^tm1a(EUCOMM)Hmgu^* allele were subsequently crossed with the ROSA26::FLPe knock in deleter line (JAX #009086, RRID:IMSR_JAX:009086) to generate *Nrf2^fl/+^* mice. These were intercrossed to obtain animals homozygous for the floxed allele. *LysM^Cre/+^* mice were purchased from Jackson labs (JAX #004781, RRID:IMSR_JAX:004781). *LysM^Cre/+^*:*Blvra^Δ/Δ^* (*Blvra^LysMΔ/Δ^*) mice were generated at GIMM by crossing *Blvra^fl/fl^* ^42^ mice with *LysM^Cre/+^:Blvra^+/fl^* mice. *LysM^Cre/+^:Nrf2^Δ/Δ^* (*Nrf2^LysMΔ/Δ^*) mice were generated at GIMM by crossing *Nrf2^fl/fl^* mice with *LysM^Cre/+^:Nrf2^+/fl^* mice. *LysM^Cre/+^:Nrf2^Δ/Δ^:Blvra^Δ/Δ^* (DKO*^LysMΔ/Δ^)* were generated at GIMM by crossing *Nrf2^+/fl^Blvra^fl/fl^* with *LysM^Cre/+^*: *Nrf2^+/fl^:Blvra^fl/fl^*. *Spic^-/-^*animals were generated at GIMM from frozen sperm from *Spic^tm1(EGFP/cre/ERT2)Wtsi^*(EUROCOMMTools) by IVF into C57BL/6J oocytes. Offspring was subsequently bred to homozygosity and animals fully characterized.

For iron deficient diet experiments, mice were fed *ad libitum* iron deficient diet or correspondent control diet (U8958A01R 00176 or U8958A01R 00177, respectively; SAFE), starting at 7 weeks of age and maintained under SPF conditions and housed at standard *vivarium* temperature (22°C) in a 12-hour light/dark cycle with free access to water.

## METHOD DETAILS

### Bone marrow-derived macrophage (BMDM) generation

Bone marrow was isolated from femora and tibiae of wild type or indicated mutant mice. Bones were flushed with Roswell Park Memorial Institute medium (RPMI, GIBCO) containing 10% heat-inactivated fetal bovine serum (FBS, GIBCO) and 1% penicillin and streptomycin (Pen/Strep,ThermoScientific). Erythrocytes were lysed by resuspending the pellet in red blood cell lysis buffer (150 mM NH4Cl; 10 mM KHCO3; 130 μM ethylenediaminetetraacetic acid (EDTA)). The reaction was stopped by adding RPMI and cells were centrifuged. Primary mouse BMDM were cultured for 6 days in RPMI supplemented with 10% FBS, 10% L929 conditioned media and 1% Pen/Strep. BMDM, at a density of 1,.5 x10^5^ cells/well in 96 well plates, 1 x 10^6^ cells/well in 12-well plates or 2 x 10^6^ cells/well in 6-well plates, were seeded into growth media overnight before cell stimulation.

### Heme preparation

Stock solutions of high purity heme (Hemin; #H651-9, Frontier Specialty Chemicals) were prepared as described^45^. Briefly, hemin was dissolved in 0.2 N NaOH using 0.2 N HCl to adjust to pH 7.4. Solutions were centrifuged at 2792*g* for 15 min at 4°C, the supernatant was filtered using a 70 µm cell strainer and stored at-80°C. Absorbance was determined using a spectrophotometer (SmartSpec 3000, Bio-rad) at λ_405_nm. Concentrations were calculated using heme extinction coefficient (EmM = 85.82 in DMSO), following the Lambert-Beer law (A_405_nm = εcl).

### Cell stimulation

BMDM were stimulated in RPMI containing 10% FBS and 1% penicillin and streptomycin, with the following PAMPs, DAMPs, inhibitors and chemicals alone or in combinations when indicated: RSL3 (125 nM, Selleckchem), ferric ammonium citrate (FAC, 200 µM, Merck), heme (50 µM, Frontier Specialty Chemicals), bilirubin (200 nM, Frontier Specialty Chemicals), lipopolysaccharide (LPS, tlrl-3pelps, 1-15ng/µL, InvivoGen), nigericin (5-10 µM, Sigma), TNF (1-100 ng/mL, Gibco), Z-VAD(OMe)-FMK (zVAD; 25 µM, Cayman Chemical), Necrostatin 2 racemate (Nec-1s; 45 mM, Selleckchem), Deferoxamine (DFO; 50 mM, Cayman Chemical), Liproxstatin-1 (Lip-1, 200 nM, Cayman Chemical).

### Real-time imaging for cell death

Cell death kinetics was monitored using a BioTek Cytation 5 (Agilent) Cell Imaging Multimode Reader, equipped with a Teledyne CMOS camera (CM3-U3-50S5M), 20x Olympus Plan Fluo 0.45 NA dry-immersion objective, 365 and 523 nm LED lines, and DAPI and PI specific filter cubes from Agilent (1225100 and 1225111, respectively. Cells were plated in 96 well plates and stained with 100nM of Hoechst for 30 min prior to treatment with the indicated stimuli. Cell death was measured by propidium iodide (PI, 0.2µg/mL, P3566, Life Technologies) incorporation following the manufacturer’s protocol. The plate was imaged in fluorescence and brightfield modes for the indicated time durations, with a time interval of 3 h. At least 3 technical replicates were performed, with 9 images being acquired per replicate per time point. Image analysis was performed using the BioTek Gen5 Software for Imaging & Microscopy (v.1.16). Briefly, Hoechst and PI channels were background flattened with a rolling-ball diameter of 21 µm to improve cell detection. Cell masks were identified by thresholding the Hoechst signal, using controls for negative staining as reference. Dead cells were identified by defining a sub-population of Hoechst-positive cells that were also PI-positive, using a positive control for cell death as reference. Cell death ratios were then calculated from the identified double-positive cells and Hoechst positive cells.

### RNA extraction and qRT-PCR

Total RNA was extracted using tripleXtractor reagent (GRISP chlorophorm, isopropanol and ethanol, according to manufacturer’s instructions. cDNA was synthesized using the Xpert cDNA Synthesis Mastermix (GRiSP), followed by qRT-PCR using the iTaq Universal SYBR Green Supermix (Bio-Rad) on a QuantStudio™ 7 Flex Real-Time PCR System (Applied Biosystems). Transcript values were calculated from the threshold cycle (Ct) of each gene using the 2^-ΔCT^ method using Ribosomal protein lateral stalk subunit P0 (*Rplp0*) as the housekeeping control gene.

### Serology and Histopathology

Blood was collected via cardiac puncture with a syringe equipped with a 21G needle and all whole blood and serological analysis was performed by an external veterinarian clinical laboratory (DNAtech; Portugal; http://www.dnatech.pt/web/). Mice were transcardially perfused *in toto* with ice-cold PBS (1X, 15 mL) and organs were harvested, fixed (10% formalin), embedded in paraffin, sectioned (3 µm) and stained with Hematoxylin & Eosin (H&E) and Prussian blue. Whole sections from formalin-fixed and H&E-stained tissues were analyzed with a DMLB2 microscope (Leica), and images were acquired with a DFC320 camera (Leica) and NanoZoomer-SQ Digital slide scanner (Hamamatsu Photonics). Histology was analyzed at the GIMM Histopathology Unit by Dr. Pedro Faísca.

### Western Blot

BMDM were lysed using NP40 extraction buffer (0,15M NaCl, 1% NP-40, 0,05M Tris, 1X protease inhibitor cocktail [(cOmplete™, Mini, EDTA-free Protease Inhibitor Cocktail; Roche) and sonicated (Branson)]. Supernatant was collected and protein was quantified via Bradford assay (BioRad, #5000006). Protein was resolved (50µg) on a 6-12% SDS-PAGE and transferred to Polyvinylidene fluoride (PVDF) membranes. Membranes were blocked for 1 hour at room temperature (5% bovine serum albumin in 1X TBS-T), washed in 1X TBS-T and incubated with primary antibodies, overnight at 4°C. The primary antibodies used were rabbit polyclonal anti-BVRA (Thermofisher scientific, #PA5-92059; 1:750), rabbit monoclonal anti-NRF2 (CST, #D1Z9C, 1:1000), rabbit polyclonal anti-β-actin (Cell Signaling Technology, #4967; 1:1000) and rabbit monoclonal Anti-Vinculin (Cell Signaling Technology, #13901;1:1000). Membranes were washed 3 times (1X TBS-T) and incubated (1 h; RT) with the peroxidase-conjugated secondary antibody (HRP conjugated goat anti-rabbit IgGH+L; Invitrogen, #31460; 1:5000). Membranes were washed 3 times (1X TBS-T) and peroxidase activity was detected using SuperSignal™ West Pico PLUS Chemiluminescent Substrate (ThermoFisher Scientific). Blots were developed using Amersham Imager 680 (GE Healthcare), equipped with a Peltier cooled Fujifilm Super CCD or using an iBright FL1500 (ThermoFisher Scientific). Densitometry analysis was performed using ImageJ software, from images without saturated pixels.

### Total iron measurements

Total liver iron levels were measured as previously described^46^, with the following modifications to detect total iron levels. Briefly, 100 μL (20 mg tissue) of tissue homogenate were hydrolyzed at 65 °C with 50 μl TCA-HCl solution (16 h) and boiled 1 h at 120 °C to release heme. Clarified samples were incubated with sodium acetate, bathophenanthroline disulfonic acid and ascorbic acid solution for 5 min at 21-23 °C, absorbance was measured at 540 nm and iron concentration was calculated using [Fe] = (((A_s_ - A_b_) × V × MW))/((e × l × t)), (A_s_: sample absorbance, A_b_: blank absorbance, V: volume, MW: molecular weight of iron, e: milimolar absorptivity of bathophenanthroline disulfonic acid, l: path length and t: weight of tissue). Measured concentrations were verified against standard samples with known iron concentrations.

### RBC injection

Whole blood was collected from C57BL/6J mice and centrifuged (400g, 10 min) to remove the buffy coat and washed three times in PBS. For PKH26 labelling, RBCs (1 x 10¹⁰) were resuspended in diluent C (1 mL), mixed with diluent C (1 mL) containing PKH-26 (4 μM; Sigma-Aldrich, St. Louis, MO, USA) and incubated in the dark at room temperature (RT) for 5 min. The reaction was stopped by addition of PBS (10 mL) containing BSA (0.5%) and FCS (2%). RBCs were washed twice, resuspended in PBS, and centrifuged (400g, 15 min) to reduce the volume to 1 mL. RBC count and PKH26 staining efficiency were assessed by flow cytometry. For mouse experiments, RBCs (2.5 x 10⁷) were injected within 1 hour of isolation into animals aged 12 to 16 weeks.

### Flow Cytometry

For all experiments, mice, aged 16 weeks, were sacrificed by CO_2_ asphyxiation, transcardially perfused *in toto* with ice-cold PBS with EDTA (1X, 20 mL).

### Spleen, lungs and liver myeloid panels

Spleens, lungs and liver were harvested and digested with RPMI media containing 1mg/mL collagenase D (for spleen), collagenase IV (for lungs) or collagenase VII (for liver) and DNase I. Cell suspensions were passed through a cell strainer (70µm, Corning) and RBCs were lyzed using ACK buffer. Cells were then incubated with Fc block (Biolegend, Cat #553142) and fixable live dead yellow dye (LIVE/DEAD™ Fixable Yellow Dead Cell Stain Kit, Invitrogen, Cat #L34959) for 20 min (4°C). Cells were washed with PBS containing 2% FBS and centrifuged (800*g*, 2 min., 4°C) and stained (25 min., 4°C) with following antibodies: For spleen, CD45 APCe780 (Invitrogen, Cat #47-0451-82), CD11b BV785 (Biolegend, Cat #101243), F4/80 BV421 (Biolegend, Cat #123132) or PE #47-0451-82), CD11b BV785 (Biolegend, Cat #101243), F4/80 BV421 (Biolegend, Cat #123132) or PE/Cy5 (Biolegend, Cat #123112), CD163 SB702 (Invitrogen, Cat #67163182) or BV421 (Biolegend, Cat #155309), CD11c BV605 (Biolegend, Cat # 117333), Ly6C PE/Cy7 (Biolegend, Cat #128018), Ly6G AF700 (Invitrogen, Cat #56-9668-82), CD64 PE/ Dazzle594 (Biolegend, Cat #139320), CD3 [Biotin (Biolegend, Cat #100243) or PE (BD Pharmingen, Cat #561809)], CD19 [Biotin (BD Biosciences, Cat #553784) or PE (BD Pharmingen, Cat #557399)], CD49b [Biotin (Biolegend, Cat #103521) or PE (BD Pharmingen, Cat #557420)], CD90.1 [Biotin (Biolegend, Cat #202510) or PE (Biolegend, Cat #202524)]; For lungs, CD45 APCe780 (Invitrogen, Cat #47-0451-82), CD11b BV785 (Biolegend, Cat #101243), F4/80 PE/Cy5 (Biolegend, Cat #123112), CD11c BV605 (Biolegend, Cat # 117333), SiglecF APC-R700 (BD Pharmingen, Cat #565183), Ly6C PE/Cy7 (Biolegend, Cat #128018), Ly6G AF700 (Invitrogen, Cat #56-9668-82), CD64 PE/ Dazzle594 (Biolegend, Cat #139320), CD3 Biotin (Biolegend, Cat #100243), CD19 Biotin (BD Pharmingen, Cat #553784), CD49b Biotin (Biolegend, Cat #103521), CD90.1 Biotin (Biolegend, Cat #202510); For liver, CD45 APCe780 (Invitrogen, Cat #47-0451-82), CD11b BV785 (Biolegend, Cat #101243), F4/80 BV421 (Biolegend, Cat #123132) or PE/Cy5 (Biolegend, Cat #123112), CD11c BV605 (Biolegend, Cat # 117333), Ly6C PE/Cy7 (Biolegend, Cat #128018), Ly6G AF700 (Invitrogen, Cat #56-9668-82), CD3 [Biotin (Biolegend, Cat #100243) or PE (BD Pharmingen, Cat #561809)], CD19 [Biotin (BD Pharmingen, Cat #553784) or PE (BD Pharmingen, Cat #557399)], CD49b [Biotin (Biolegend, Cat #103521) or PE (BD Pharmingen, Cat #557420)], CD90.1 [Biotin (Biolegend, Cat #202510) or PE (Biolegend, Cat #202524)]. Cells were stained with PE/Fire 700 streptavidin (Biolegend, Cat #405174) diluted in PBS containing 2%FBS. Samples were resuspended in PBS containing 2%FBS and acquired on the BD LSR Fortessa X-20 cell analyzer (BD Biosciences) or on the Cytek Aurora analyzer (Cytek).

### Peritoneal lavage fluid (PLF)

Mice were sacrificed and injected 5 mL cold sterile PBS. After this, the abdomen was massaged and the injected fluid was recovered. The recovered PLF was then centrifuged (300*g*, 5 min., 4°C) and RBCs were lyzed using ACK buffer. Cells were then incubated with Fc block (Biolegend, Cat #553142) and fixable live dead yellow dye (LIVE/DEAD™ Fixable Yellow Dead Cell Stain Kit, Invitrogen, Cat #L34959) for 20 min (4°C). Cells were washed with PBS containing 2% FBS and centrifuged (800*g*, 2 min.,4°C) and stained for 25 min at 4°C with following antibodies: CD45 APCe780 (Invitrogen, Cat #47-0451-82), CD11b BV785 (Biolegend, Cat #101243), F4/80 PE/Cy5 (Biolegend, Cat #123112), CD11c BV605 (Biolegend, Cat # 117333), Ly6C PerCP/Cy5.5 (Biolegend, Cat #128012), Ly6G AF700 (Invitrogen, Cat #56-9668-82), MHC-II eF506 (Invitrogen, Cat #69-5321-82), TIM-4 PE/Vio615 (Miltenyi Biotec S. L, Cat # 130-116-638), CD3 Biotin (Biolegend, Cat #100243), CD19 Biotin (BD Pharmingen, Cat #553784), CD49b Biotin (Biolegend, Cat #103521), CD90.1 Biotin (Biolegend, Cat #202510). Cells were stained with PE/Fire 700 streptavidin (Biolegend, Cat #405174) diluted in PBS containing 2%FBS. Samples were resuspended in PBS containing 2%FBS and acquired on the Cytek Aurora analyzer (Cytek).

### BMDM ROS and lipid peroxidation measurement analysis

BMDM were stimulated in RPMI 1640 with Glutamax containing 10% FBS and 1% Pen/Strep, with the 125-500 nM RSL3 (Selleckchem) and 200 nM ferric ammonium citrate (FAC, Merck) for 6h. Cells were then resuspended and stained with 2′,7′-dichlorodihydrofluorescein diacetate (H2DCFDA, AP Bio, #C263) for reactive oxygen species (ROS) measurement and Liperfluo ( Dojindo, #B3932) for lipid peroxidation for 30 min at 37°C. Samples were resuspended in PBS containing 2%FBS and acquired on the Cytek Aurora analyzer (Cytek).

### Bulk RNA sequencing

Full-length cDNAs were generated following the SMART-Seq2 protocol described^47^. After quality control using Fragment Analyzer (Agilent Technologies), library preparation including cDNA ‘tagmentation’, PCR-mediated adaptor addition and amplification of the adapted libraries was done following the Nextera library preparation protocol (Nextera XT DNA Library Preparation kit, Illumina), as previously described^48^. Libraries were confirmed by Fragment Analyzer (Agilent Technologies) and then sequenced (NextSeq2000, Illumina) using 100 SE P2. Sequence information was extracted in FastQ format, using Illumina DRAGEN FASTQ Generation v3.8.4. Library preparation and sequencing were optimized and performed by Genomics Platform at the GIMM. FastQ reads were trimmed using fastp (v0.23.4) and aligned to GRCm39 (Ensembl release 111) with STAR (v2.7.11b). Reads were summarized at the gene level using featureCounts (v2.0.2). Differential expression analysis was performed with DESeq2 (v1.46.0) in R (v4.4.2), using the design ∼genotype + treatment + genotype:treatment (genotype: Spic-KO or WT; treatment: heme or control; N = 3 per group). Genes with mean raw counts ≤5 were excluded (14,187 retained). DEGs were defined by adjusted p-value < 0.05 and |log₂FC| > 0.585, using unshrunken estimates. PCA was performed on variance-stabilized values (blind = TRUE). Heatmaps were generated with pheatmap (v1.0.12) using Z-score scaling and complete-linkage clustering. Volcano plots were generated with ggplot2 (v3.5.2). ORA was performed separately on up-and downregulated DEGs using Fisher’s exact tests with Benjamini-Hochberg correction, against gene sets curated from GO, KEGG, Reactome, and transcription factor targets.

### Single cell RNA sequencing

Spleen cells were isolated as described above, from animals aged 14 weeks. The homogeneous cell suspension of splenocytes was subjected to sorting with the Cytek Aurora CS cell sorter (Cytek) using a 100 µm nozzle. Single, live, CD45^+^, CD64^+^ cells were selected for sorting and collected in 100 µL FACS buffer. Single-cell experiments were performed using 10x Genomics technology. Following cell sorting, suspensions were adjusted to approximately 1,500 cells/µL and processed using the Chromium Controller with the Chromium GEM-X Single Cell 3’ Reagent Kit v4 (Dual Index) (PN-1000691, 10x Genomics, Pleasanton, CA, USA), following the manufacturer’s protocol. Cells were loaded at 30,000 cells/sample for library preparation and sequencing. cDNA integrity and concentration, as well as library concentration and size distribution, were assessed using a Fragment Analyzer (Agilent Technologies). Quantification was performed using a high-sensitivity dsDNA assay (Qubit, Thermo Fisher Scientific). For final library preparation, amplified cDNA was fragmented and size-selected using SPRIselect magnetic beads (Beckman Coulter) to obtain fragments of 300-800 bp. After adaptor ligation and sample indexing, fragment size and concentration were re-assessed, and libraries were pooled in equimolar amounts. Sequencing was performed at ∼30 million reads/cell with 28 bp R1 and 90-cycle R2 paired-end reads on an Illumina NextSeq 2000 system. To minimize technical variation, N=3 *Ctrl^fl/fl^*and N=4 *DKO^LysMΔ/Δ^* animals were processed and sequenced in parallel.

### Single-cell data alignment, read processing, data analysis and annotation

Raw sequencing data were demultiplexed and aligned to the *Mus musculus* reference genome (GRCm38/mm10) using Cell Ranger 7.1.0, with intronic reads included. The resulting count matrices were processed using Scanpy (v1.8.2). Low-quality cells were excluded based on the following thresholds: 600-8,000 genes per cell, 500-150,000 UMIs per cell, and mitochondrial gene content <15%. Non-informative genes, including *Malat1*, mitochondrial-encoded, ribosomal, and hemoglobin genes, were removed prior to downstream analysis. Doublets were identified and removed using Scrublet with a doublet score threshold of 0.22. Data were normalized, log-transformed, and highly variable genes (n = 3,000) were selected. PCA was performed, and UMAP was computed for two-dimensional visualization. Clusters were identified and cell types were manually annotated based on the top 20 highly expressed genes per cluster. Low-level carry-over of adaptive immune cells (B and T cells) into the CD64⁺ fraction was handled computationally by stringent quality control, doublet removal, and marker-guided annotation; these populations were excluded from all macrophage-centric analyses. Sub-clustering of the macrophage compartment was performed to resolve developmental intermediates along the monocyte-to-RPM axis based on *Spic* expression. A numerically minor NF-κB-activated RPM cluster identified during sub-clustering was excluded from downstream comparisons owing to its small size. B cells resolved as a distinct cluster during sub-clustering were likewise excluded. Genotype-dependent transcriptional differences within each RPM-lineage cluster were assessed by pseudobulk differential expression: raw counts were aggregated per sample per cluster and compared between *Ctrl^fl/fl^* and *DKO^LysMΔ/Δ^* using DESeq2 (PyDESeq2), with Benjamini-Hochberg correction (adjusted α < 0.05). A ferroptosis suppression module score was computed per cell using scanpy.tl.score_geneS8 with a gene set defined a priori from FerrDb v2^30^ and KEGG pathway mmu04216, comprising *Gpx4*, *Slc7a11*, *Gclm*, *Gclc*, *Gss*, *Gsr, Txnrd1, Hmox1, Nqo1, Blvra, Fth1, Nfs1, Cisd1, and Aifm2*. This gene set was defined independently of differential expression results in the dataset. Per-sample scores were compared between genotypes using the two-tailed Mann-Whitney U test with Benjamini-Hochberg correction; significance thresholds: \**p* < 0.05, \*\**p* < 0.01, \*\*\**p* < 0.001.

### Metabolomics

BMDM were stimulated with 50 µM heme for 24h. Metabolites were extracted with 1 mL of ice-cold extraction solvent composed of 80% methanol, acetonitrile and water (4:4:2) containing 1.5 µM metabolomics amino acid mix (Cambridge Isotope Laboratories, cat. no. MSK-A2-1.2). Samples were transferred to 1.5 mL Eppendorf tubes, centrifuged at 20,000g for 20 min at 4°C. All the supernatant was transferred to a new tube, dried down in a SpeedVac (Thermo Scientific) for 2 h and stored at-80°C until analysis. The dried extracts were then resuspended in 40 μL of 60:40 acetonitrile:water, vortexed and incubated on ice for 20 min. After this incubation period, 10 μL of 100% methanol was added and the extracts were centrifuged at 20,000g for 20 min at 4°C to remove any particulates. LC-MS analysis was performed on a Thermo Scientific Vanquish Flex UHPLC system coupled to an Orbitrap Exploris 240 mass spectrometer (Thermo Scientific). Chromatographic separation was achieved using a XBridge BEH Amide column (2.5 µm; 2.1×100 mm, Waters). The mobile phases consisted of 10 mM ammonium acetate in water with 0.1% ammonium hydroxide as mobile phase A (MPA) and 10 mM ammonium acetate in 90% acetonitrile with 0.1% ammonium hydroxide as mobile phase B (MPB). InfinityLab Deactivator Additive (Agilent Technologies) was added to both MPA and MPB at 5 µM. Both mobile phases were sonicated overnight to prevent precipitation. The column temperature was maintained at 40°C, with an injection volume of 2 µL and a flow rate of 0.4 mL/min. The chromatographic gradient was as follows: 0 min, 100% B; 12.5 min, 90% B; 19 min, 60% B; 20 min, 60% B; 20.5 min, 100% B; and 25 min, 100% B. MS data acquisition was performed in both positive and negative ionization modes, with spray voltages of 3,500 V and 2,500 V, respectively. Other parameters were sheath gas, 50 arb; auxiliary gas, 10 arb; sweep gas, 1 arb; ion transfer tube temperature, 325°C; vaporizer temperature, 350°C; AGC target, in standard mode; and maximum injection time in auto mode. Data were acquired in full-scan mode (m/z 70-1,000) at ×60,000 resolution using the Xcalibur software package (Thermo Scientific). Peak identification was performed by matching accurate mass (±5 ppm) and retention time to reference standards in an in-house library. For metabolites S7P, E4P, R5P, and N-CA, retention time was not yet confirmed with a reference standard. Data analysis was performed with Skyline v26.1.

## QUANTIFICATION AND STATISTICAL ANALYSIS

For non-sequencing experiments, statistically significant differences between two groups were assessed using a two-tailed unpaired Mann-Whitney test. Comparisons between more than two groups were assessed using two-way ANOVA with Tukey’s multiple comparison test or two-tailed Mann-Whitney test. These analyses were performed using GraphPad Prism 8.

For bulk RNA-seq, differential expression was performed with DESeq2 using the Wald test. DEGs were defined by Benjamini-Hochberg adjusted p-value < 0.05 and |log₂FC| > 0.585. Over-representation analysis was performed using one-sided Fisher’s exact tests with Benjamini-Hochberg correction.

For scRNA-seq, genotype-dependent transcriptional differences within RPM-lineage clusters were assessed by pseudobulk differential expression using PyDESeq2, with Benjamini-Hochberg correction (adjusted α < 0.05). Ferroptosis suppression module scores were compared between genotypes using the two-tailed Mann-Whitney U test with Benjamini-Hochberg correction.

Differences were considered statistically significant at *p* < 0.05. NS: not significant, *p* > 0.05; \**p* < 0.05; \*\**p* < 0.01; \*\*\**p* < 0.001; \*\*\*\**p* < 0.0001.

---

**Figure S1.**
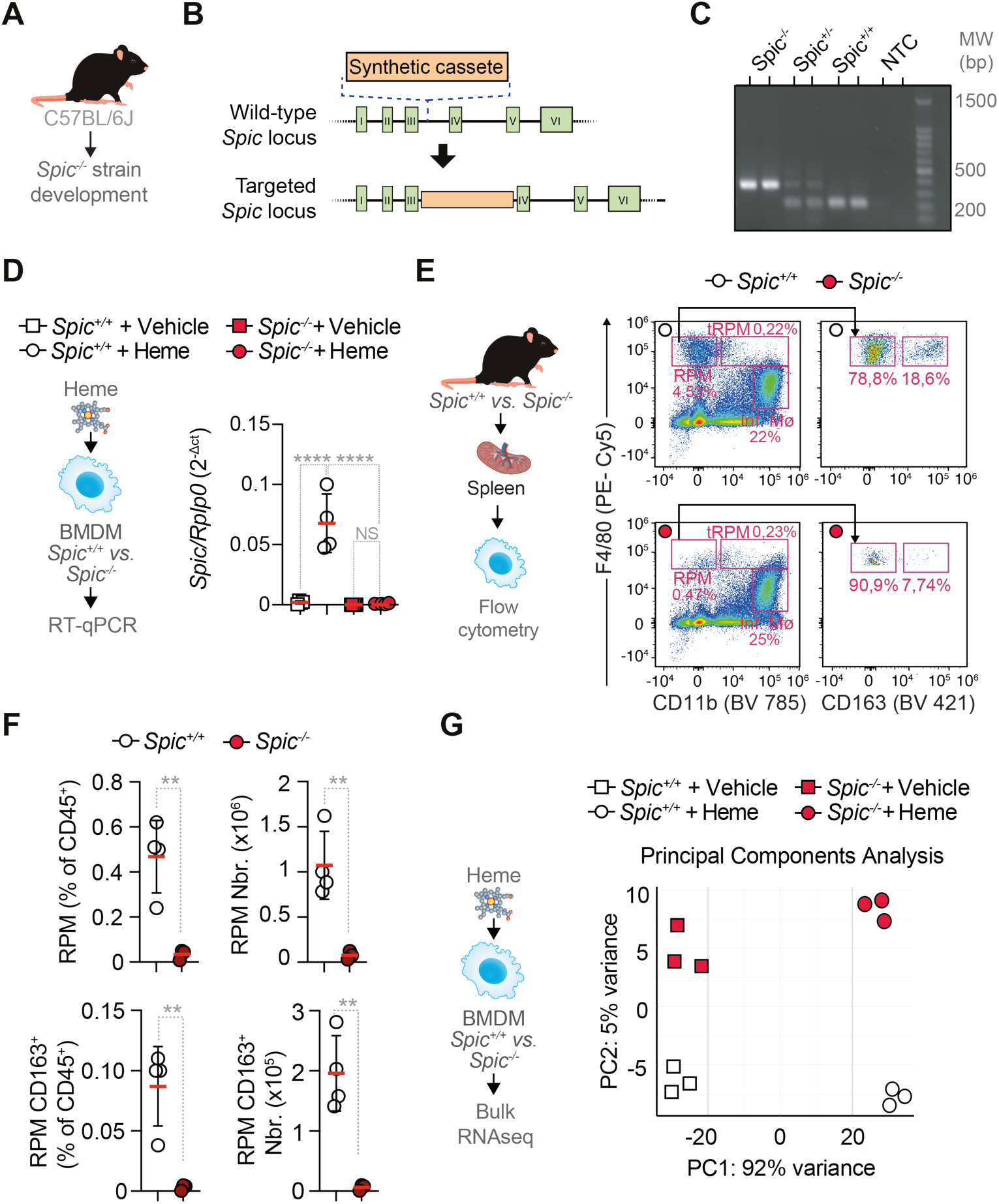
Generation of *Spic*^−/−^ mice on a C57BL/6J background (#. Figure 1**) (A,B)** Schematic representation of the targeting strategy for insertion of a synthetic cassette into the *Spic* locus between exons III and IV, disrupting exons IV–VI. **(C)** Representative genotyping PCR confirming correct targeting of the *Spic* locus in *Spic^-/-^*, *Spic^+/-^*, *Spic^+/+^* and no-template control (NTC) samples. Molecular weight (MW) markers are indicated in base pairs (bp). **(D)** RT-qPCR quantification of *Spic* expression in *Spic*^+/+^ and *Spic*^−/−^ BMDM treated with vehicle or heme. **(E)** Representative flow cytometry dot plots of splenic myeloid populations in *Spic*^+/+^ and *Spic*^−/−^ mice, gated by F4/80 and CD11b expression to identify RPM, tRPM, and inflammatory monocytes/macrophages (Inf. Mø). CD163 expression within the RPM gate is shown in the right panels. **(F)** Frequency (% of CD45^+^) and absolute numbers of RPM and CD163^+^ RPM in *Spic*^+/+^ and *Spic*^−/−^ spleens. **(G)** Principal component analysis (PCA) of bulk RNA-seq data from *Spic*^+/+^ and *Spic*^−/−^ BMDM treated with vehicle or heme. Data are presented as individual data points with mean ± SD. Data in (D) are from at least two independent experiments, n=3–4 replicates *per* condition. Data in (E) and (F) are representative of at least two independent experiments, n=3–4 mice *per* condition. Data in (G) are from one experiment, n=3 technical replicates *per* condition. NS, not significant; **p < 0.01, ****p < 0.0001 by two-tailed unpaired t-test or two-way ANOVA with Tukey’s multiple comparisons test.

**Figure S2.**
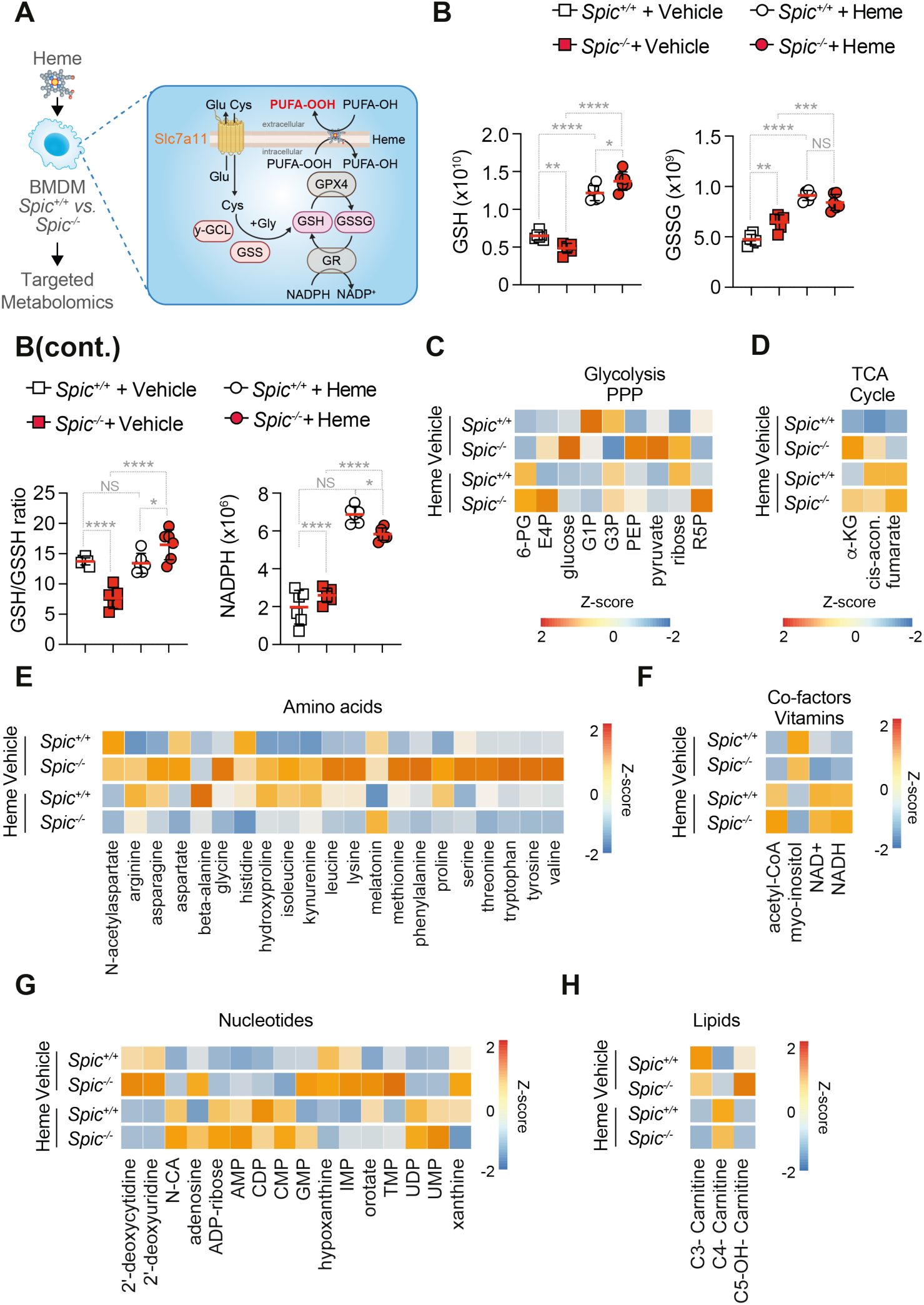
SPI-C regulates the antioxidant metabolic response to heme in macrophages (#. Figure 1). **(A)** Schematic representation of *Spic*^+/+^ and *Spic*^−/−^ BMDM treated with heme or vehicle and subjected to targeted metabolomics analysis, focusing on glutathione biosynthetic and recycling pathway. **(B-G) (B)** Quantification of reduced glutathione (GSH), oxidized glutathione (GSSG), GSH/GSSG ratio, and NADPH in *Spic*^+/+^ and *Spic*^−/−^ BMDM treated with vehicle or heme. (C-H) Heatmaps of metabolite Z-scores in *Spic*^+/+^ and *Spic*^−/−^ BMDM under vehicle and heme conditions, grouped by metabolic pathway: glycolysis and pentose phosphate pathway (PPP) metabolites **(C)**, TCA cycle intermediates **(D)**, amino acids **(E)**, co-factors and vitamins **(F)**, nucleotides **(G)**, and lipids/ acylcarnitines **(H)**. Data in **(B)** are presented as individual data points with mean ± SD, representative of one experiment, n=4 replicates *per* condition. NS, not significant; *p < 0.05, **p < 0.01, ****p < 0.0001 by two-way ANOVA with Tukey’s multiple comparisons test.

**Figure S3.**
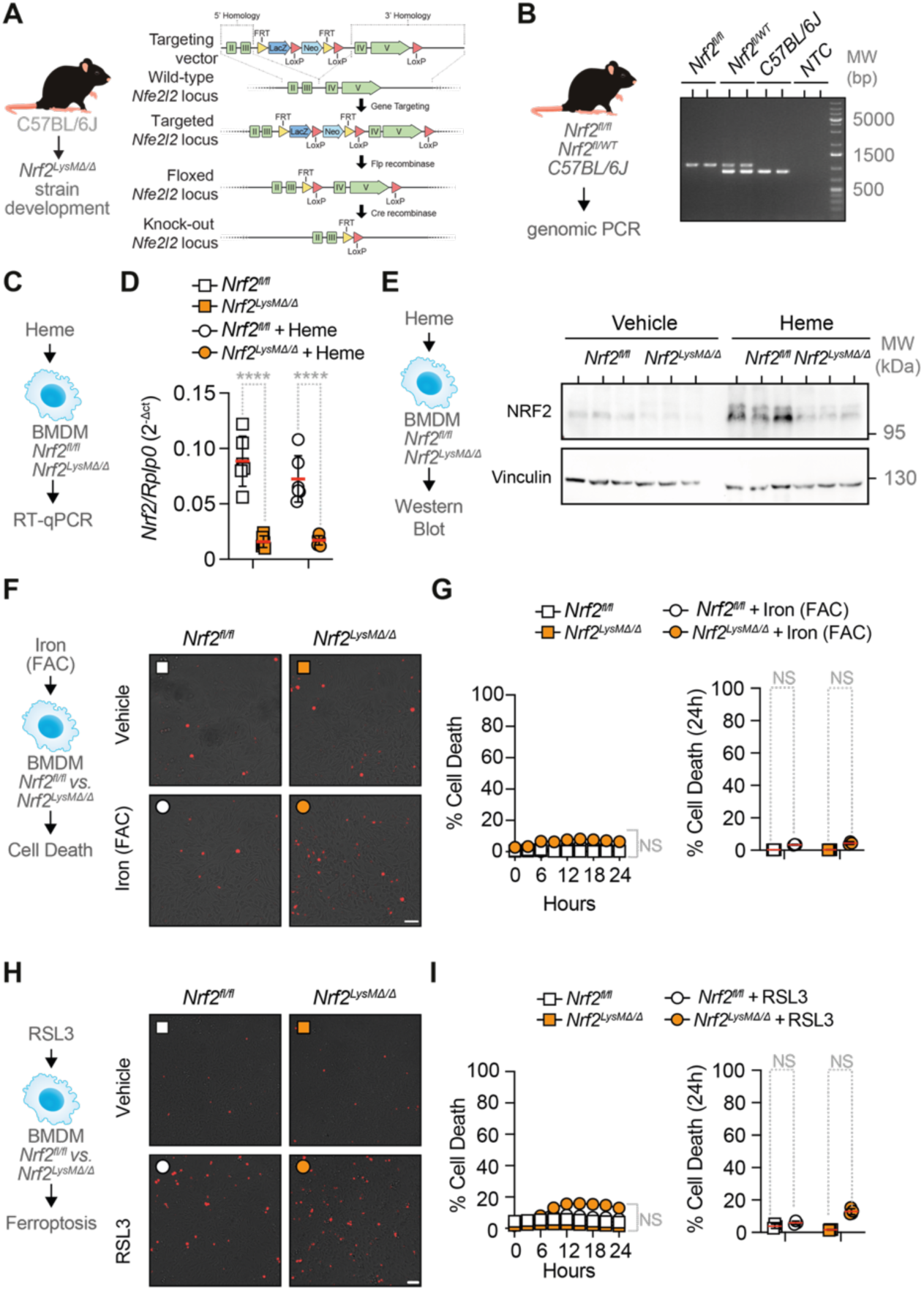
**Generation and validation of *Nrf2^LysMΔ/Δ^* mice; NRF2 deletion alone does not sensitize BMDM to ferroptosis (#**Figure 2**). (A)** Schematic representation of the strategy for the generation of *Nrf2^LysMΔ/Δ^*mice, showing the targeting vector, wild-type, targeted and floxed *Nrf2* (*Nfe2l2*) locus following Flp recombinase-mediated excision, and knock-out *Nfe2l2* locus following Cre recombinase-mediated deletion. **(B)** Representative genomic PCR confirming correct targeting of the *Nfe2l2* locus in *Nrf2^fl/fl^*, *Nrf2^fllWT^*, C57BL/6J, and NTC samples. Molecular weight (MW) markers are indicated in base pairs (bp). **(C, D)** RT-qPCR quantification of *Nrf2* expression in *Nrf2^fl/fl^* and *Nrf2^LysMΔ/Δ^*BMDM treated with heme. **(E)** Experimental schematic and representative western blot of NRF2 (∼100 kDa) and Vinculin (∼130 kDa) protein levels in *Nrf2^fl/fl^* and *Nrf2^LysMΔ/Δ^* BMDM treated with heme. **(F,G)** Microscopy images **(F)** and quantification **(G)** of cell death (red, PI-positive cells) in *Nrf2^fl/fl^* and *Nrf2^LysMΔ/Δ^* BMDM treated with iron (FAC; 200 µM for 24h). **(H and I)** Representative microscopy images **(H)** and quantification **(I)** of cell death in *Nrf2^fl/fl^*and *Nrf2^LysMΔ/Δ^* BMDM treated with RSL3 (125 nM for 24h). Data in (B) are from at least three independent experiments, n=2 technical replicates *per* condition. Data in (D,E) are from one independent experiment, n=3 technical replicates *per* condition. Data in (G) and (I) are presented as individual data points showing mean ± SEM; n=3 technical replicates representative of at least two independent experiments. Scale bars in (F) and (H) = 50 μm. NS, not significant; ****p < 0.0001 NS, not significant by two-way ANOVA with Tukey’s multiple comparisons test.

**Figure S4.**
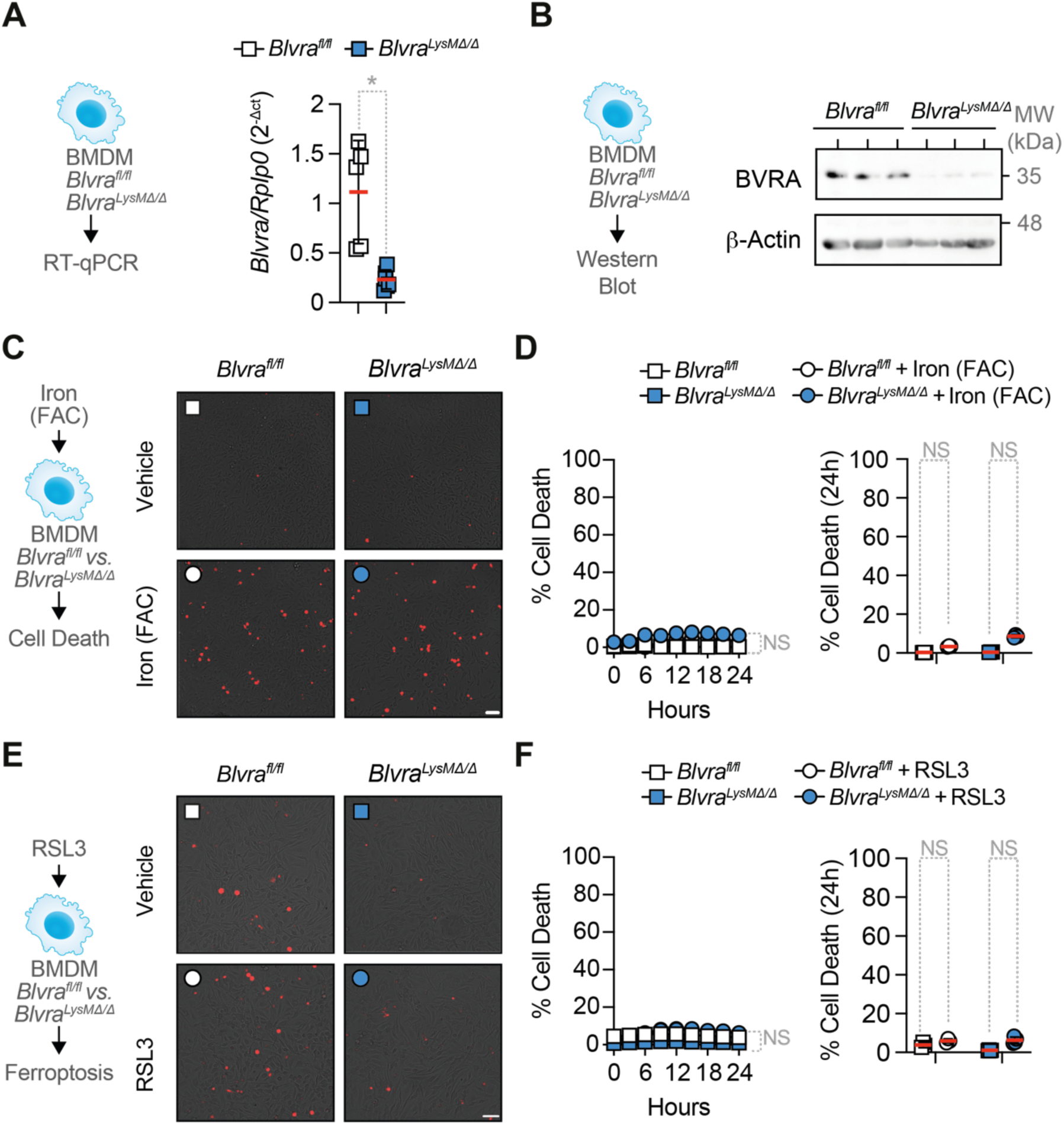
Generation and validation of *Blvra^LysMΔ/Δ^* mice; BVRA deletion alone does not sensitize BMDM to ferroptosis (#Figure 2). (A) RT-qPCR quantification of *Blvra* expression in *Blvra^fl/fl^*and *Blvra^LysMΔ/Δ^* BMDM. **(B)** Representative western blot of BVRA (∼35 kDa) and β-Actin (∼48 kDa) protein levels in *Blvra^fl/fl^*and *Blvra^LysMΔ/Δ^* BMDM. **(C,D)** Representative microscopy images **(C)** and quantification **(D)** of cell death (red, PI-positive cells) in *Blvra^fl/fl^* and *Blvra^LysMΔ/Δ^*BMDM treated with iron (FAC; 200 µM for 24h). **(E and F)** Representative microscopy images **(E)** and quantification **(F)** of cell death in *Blvra^fl/fl^*and *Blvra^LysMΔ/Δ^* BMDM treated with RSL3 (125 nM for 24h). Data in (A) is representative of two independent experiments, n=5 replicates *per* condition; (B) from one experiment, n=3 technical replicates *per* condition; (D) and (F) are presented as individual data points showing mean ± SEM; n=3 replicates representative of two independent experiments. Scale bars in (C) and (E) = 50 μm. NS, not significant; *p < 0.05 by two-tailed unpaired t-test (A) or two-way ANOVA with Tukey’s multiple comparisons test (D, F).

**Figure S5.**
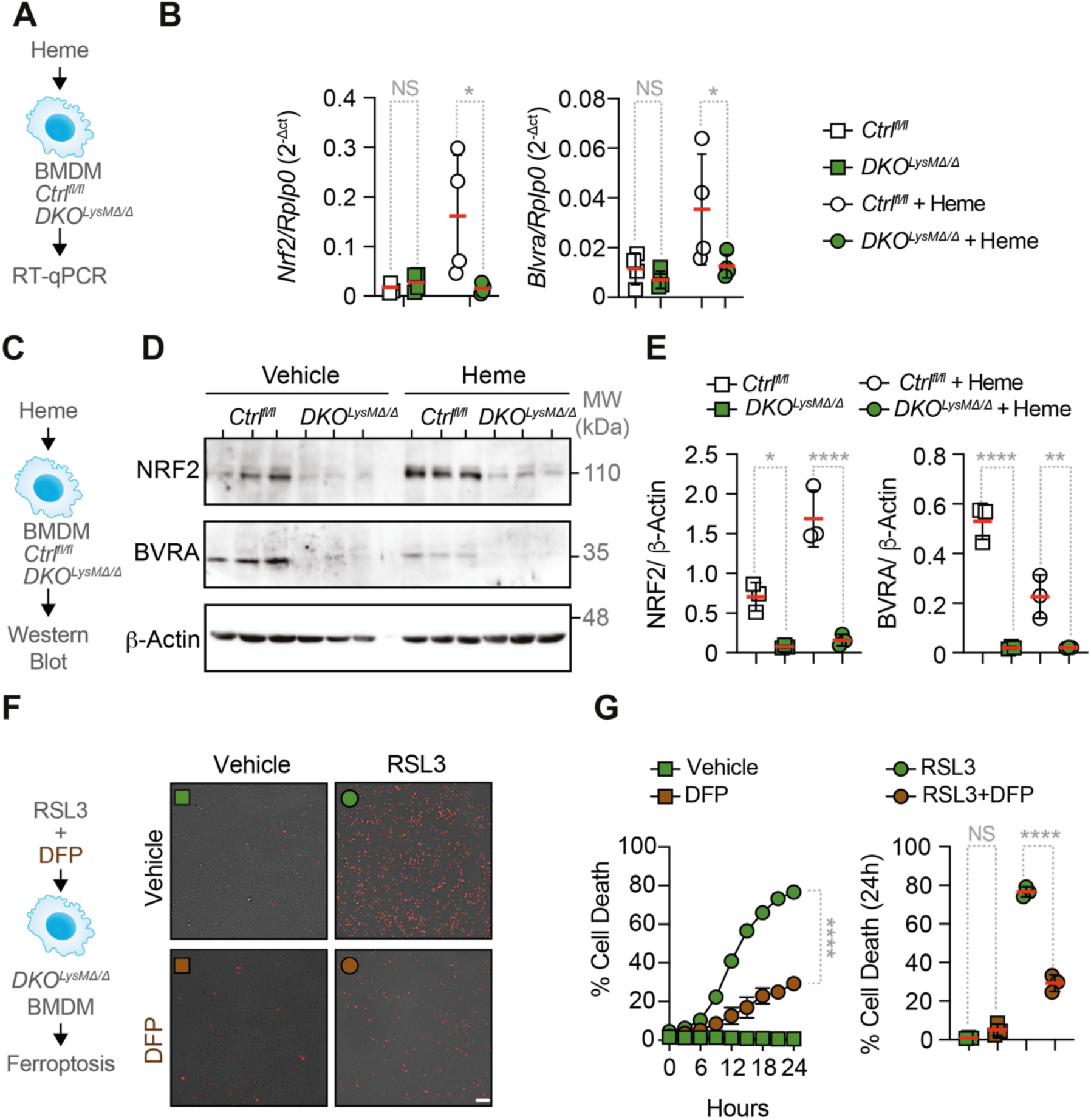
Validation of *Nrf2* and *Blvra* deletion and iron chelation rescue of ferroptosis (#. Figure 2**). (A,B)** RT-qPCR quantification of *Nrf2* and *Blvra* mRNA expression in *Ctrl^fl/fl^* and *DKO^LysMΔ/Δ^*BMDM treated with heme *vs.* vehicle. **(C, D)** Representative western blots showing NRF2 (∼110 kDa), BVRA (∼33 kDa), and β-Actin as loading control (∼42 kDa) in *Ctrl^fl/fl^* and *DKO^LysMΔ/Δ^* BMDM treated with heme *vs.* vehicle. **(E)** Quantification of NRF2 and BVRA relative to β-Actin after heme stimulation. **(F,G)** Representative microscopy images (F) and quantification (G) of cell death (red, PI-positive cells) in *Ctrl^fl/fl^* and *DKO^LysMΔ/Δ^* BMDM treated with RSL3 in the presence or absence of the iron chelator deferiprone (DFP). Scale bars, 50 μm. Data are shown as individual data points with mean ± SEM; n=3 replicates representative of two independent experiments. NS, not significant; *p < 0.05, **p < 0.01, ****p < 0.0001 by two-way ANOVA with Tukey’s multiple comparisons test..

**Figure S6.**
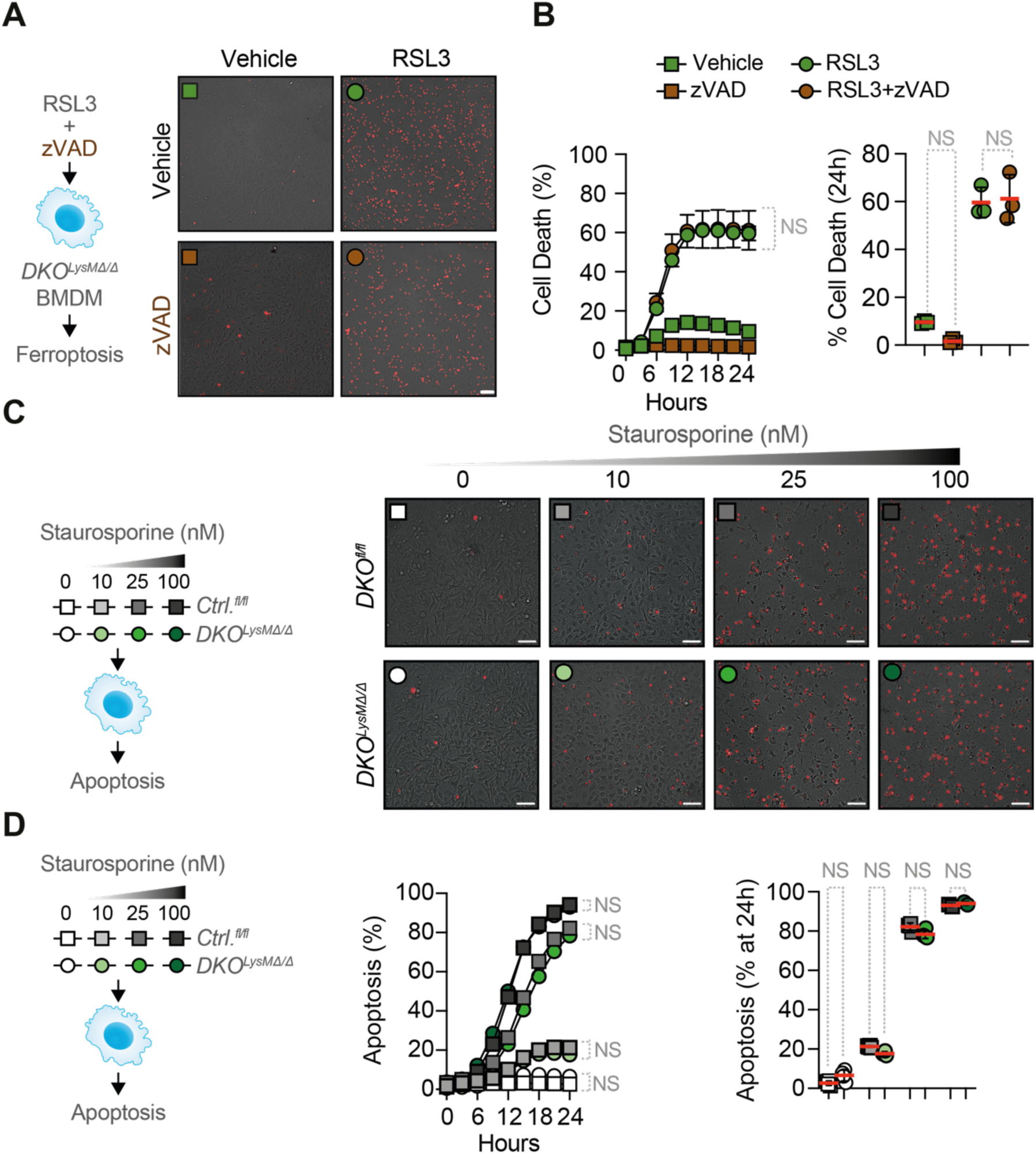
NRF2 and BVRA deficient macrophages are not sensitized to apoptosis (#. Figure 2**). (A,B)** Representative microscopy images (**A**) and quantification (**B**) of cell death (red, PI-positive cells) in *DKO^LysMΔ/Δ^* BMDM treated with RSL3 (125 nM for 24h) in the presence or absence of the pan-caspase inhibitor zVAD (25 µM for 24h). **(C and D)** Representative microscopy images (**C**) and quantification (**D**) of cell death in *Ctrl^fl/fl^* and *DKO^LysMΔ/Δ^* BMDM treated with increasing concentrations of staurosporine. Scale bars, 50 μm. Data are shown as individual data points with mean ± SEM; n=3 replicates representative of two independent experiments. NS, not significant by two-way ANOVA with Tukey’s multiple comparisons test.

**Figure S7.**
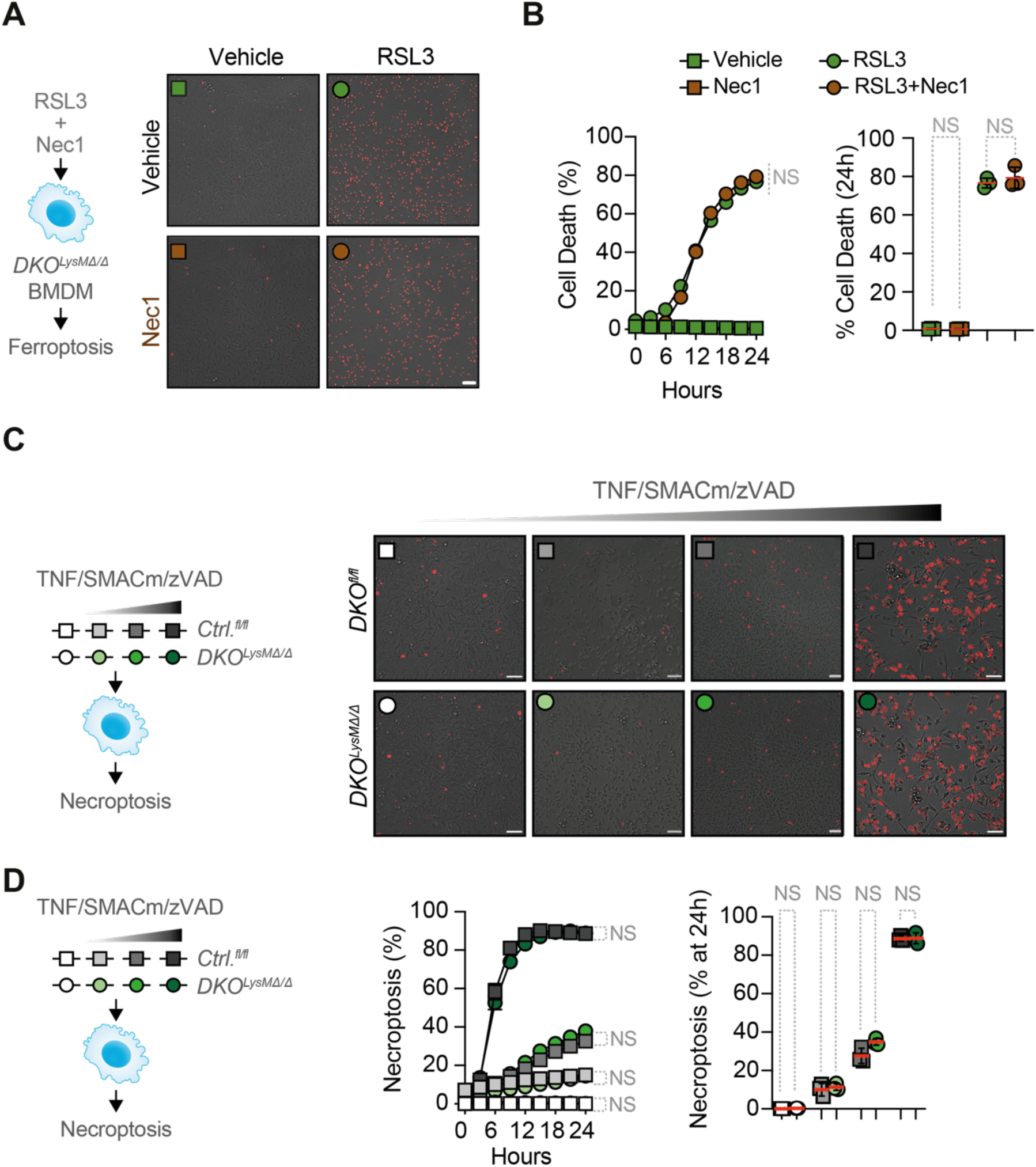
NRF2 and BVRA deficient macrophages are not sensitized to necroptosis (#. Figure 2**). (A,B)** Representative microscopy images (**A**) and quantification (**B**) of cell death (red, PI-positive cells) in *DKO^LysMΔ/Δ^* BMDM treated with RSL3 (125 nM for 24h) in the presence or absence of the necroptosis inhibitor necrostatin-1 (Nec1) (45 µM for 24h). **(C, D)** Representative microscopy images (**C**) and quantification (**D**) of cell death in *Ctrl^fl/fl^* and *DKO^LysMΔ/Δ^* BMDM treated with increasing concentrations of TNF/SMACm/zVAD (necroptosis inducer) (TNF, 1, 20, 100 ng/mL; SMACm, 100nM; zVAD, 25 µM for 24h). Scale bars, 50 μm. Data shown as individual data points with mean ± SEM; n=3 replicates representative of two independent. NS, not significant by two-way ANOVA with Tukey’s multiple comparisons test.

**Figure S8.**
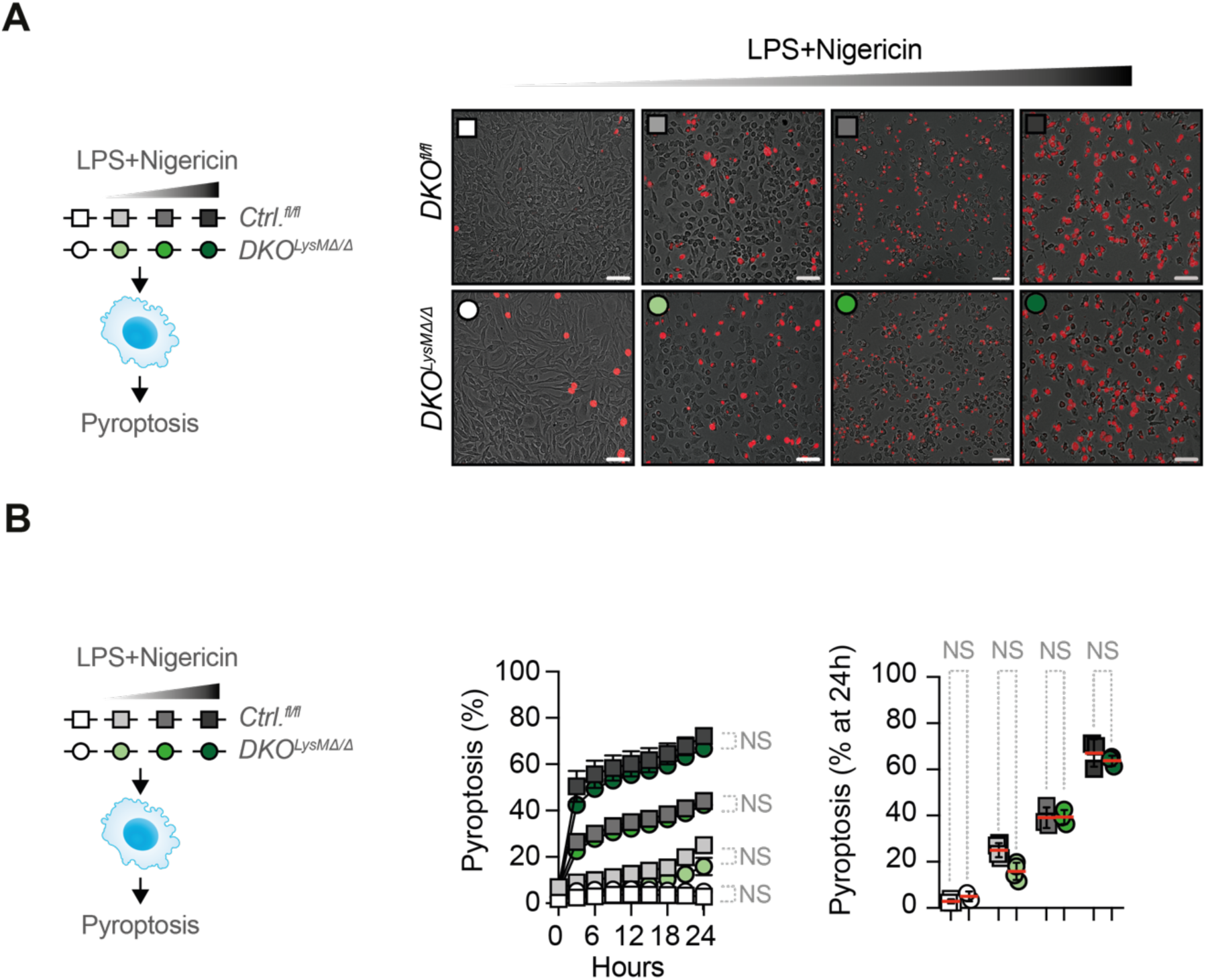
NRF2 and BVRA deficient macrophages are not sensitized to pyroptosis (#. Figure 2**). (A)** Representative microscopy images of cell death (red, PI-positive cells) in *Ctrl^fl/fl^* and *DKO^LysMΔ/Δ^*BMDM treated with increasing concentrations of LPS+Nigericin (pyroptosis inducer) (LPS, 1.5, 7.5, 15 ng/mL; Nigericin, 5µM, 10µM for 24h). **(B)** Quantification of cell death in *Ctrl^fl/fl^* and *DKO^LysMΔ/Δ^* BMDM treated with increasing concentrations of LPS+Nigericin. Scale bars, 50 μm. Data are shown as individual data points with mean ± SEM; n=3 replicates representative of two independent experiments. NS, not significant by two-way ANOVA with Tukey’s multiple comparisons test.

**Figure S9.**
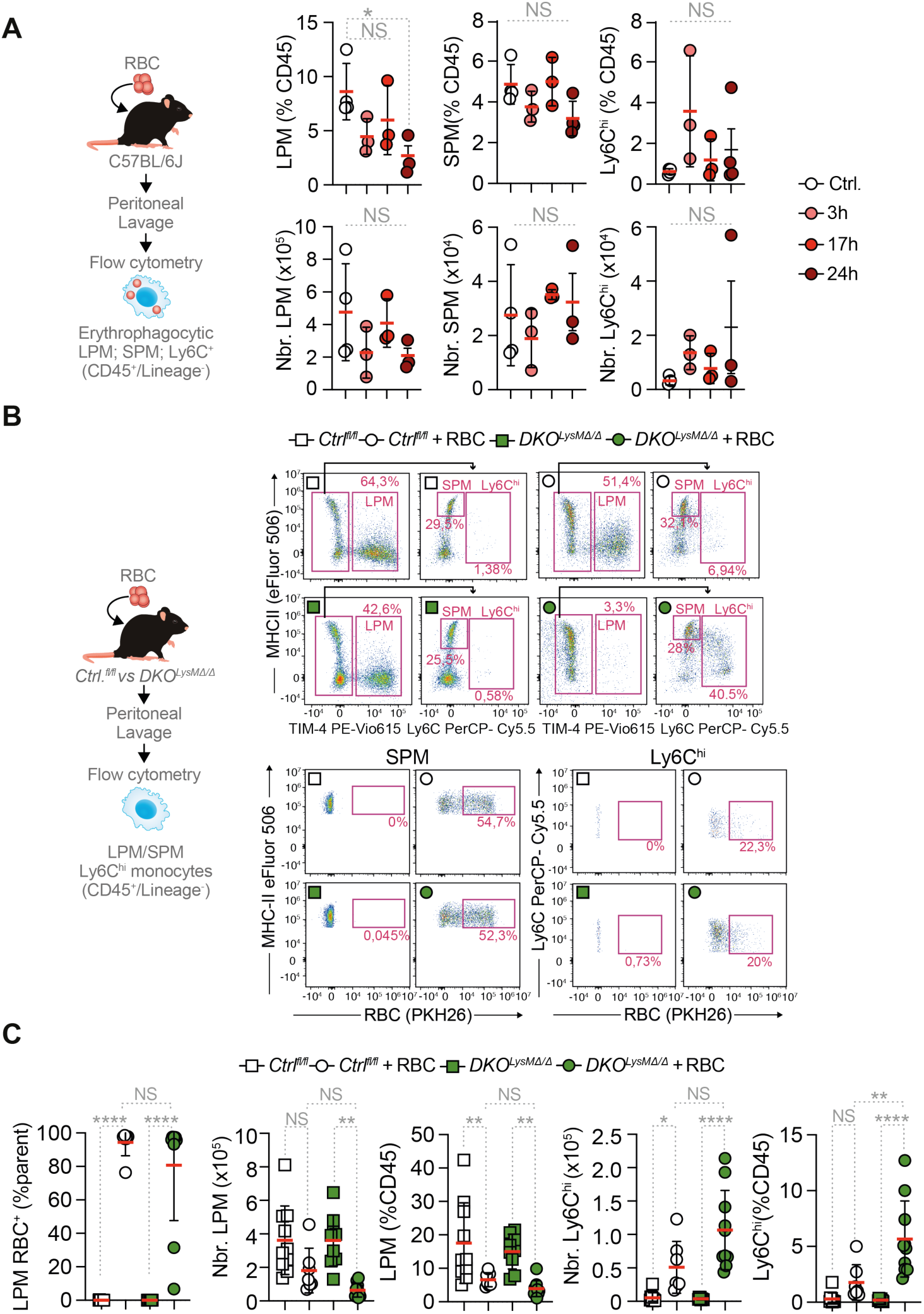
**Characterization of peritoneal erythrophagocytic populations following RBC injection (#**Figure 4**). (A)** Experimental scheme and quantification of erythrophagocytic activity (PKH26⁺) across peritoneal myeloid populations (LPM, SPM, Ly6C⁺ monocytes; CD45⁺/Lineage⁻) in C57BL/6J mice at different time points following intraperitoneal injection of PKH26-labelled RBC. **(B)** Representative flow cytometry plots showing gating strategy of peritoneal LPM, SPM and Ly6C^hi^ monocytes and respective PKH26 uptake in *Ctrl^fl/fl^* and *DKO^LysMΔ/Δ^* mice before and after RBC injection. **(C)** Quantification of erythrophagocytic activity (PKH26⁺) across peritoneal myeloid populations in *Ctrl^fl/fl^* and *DKO^LysMΔ/Δ^*mice. Data in (A) is from one experiment, n=3 mice *per* condition. Data in (C) is pooled from three independent experiments, n=7-10 mice *per* condition. NS, not significant; *p < 0.05, **p < 0.01, ****p < 0.0001 by two-way ANOVA with Tukey’s multiple comparisons test. Data are shown as individual data points with mean ± SD.

**Figure S10.**
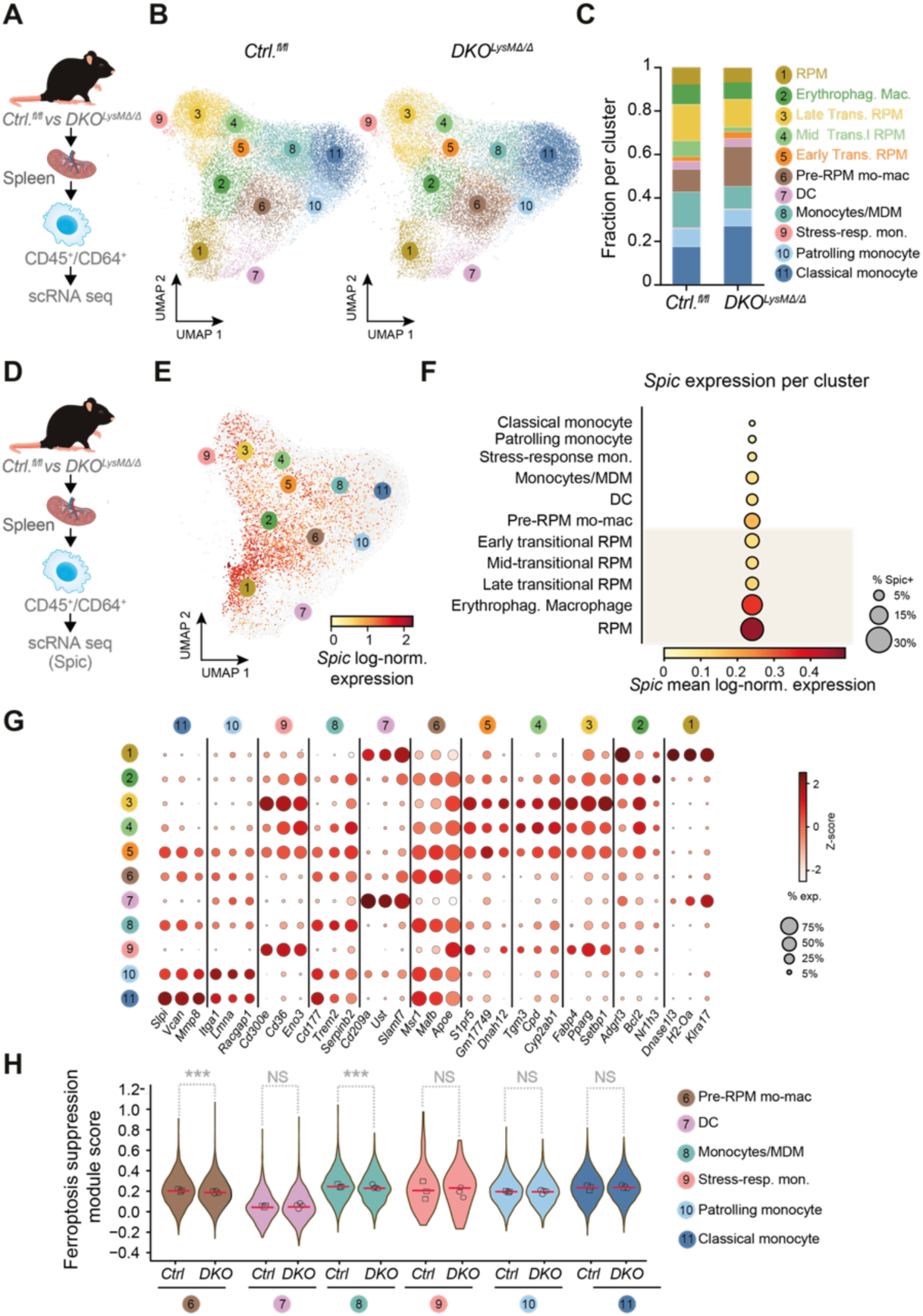
**Single-cell RNA sequencing characterization of splenic macrophages from *Ctrl^fl/fl^*and *DKO^LysMΔ/Δ^* mice (#**Figure 5). **(A)** Schematic representation of splenocytes from *Ctrl^fl/fl^*and *DKO^LysMΔ/Δ^* mice were sorted for CD45^+^CD64^+^ cells and subjected to single-cell RNA sequencing (scRNA-seq). **(B)** UMAP visualization of scRNA-seq data *Ctrl^fl/fl^*(left, n=18,972) and *DKO^LysMΔ/Δ^* (right, n=18,767) mice. Eleven transcriptionally distinct cell clusters are annotated: red pulp macrophages (RPM), erythrophagocytic macrophages, late transitional RPM, mid-transitional RPM, early transitional RPM, pre-RPM monocyte-derived macrophages (mo-mac), dendritic cells (DC), monocytes/MDM, stress-responsive monocytes, patrolling monocytes, and classical monocytes. **(C)** Stacked bar chart showing the fraction of cells per cluster in *Ctrl^fl/fl^* and *DKO^LysMΔ/Δ^* samples. **(D)** *Spic* expression analysis in CD45^+^CD64^+^ splenocytes sorted from *Ctrl^fl/fl^*and *DKO^LysMΔ/Δ^* mice. **(E)** UMAP visualization of *Spic* log-normalized expression across all sequenced cells. **(F)** Dot plot showing mean *Spic* log-normalized expression (x-axis) and percentage of *Spic*-expressing cells (dot size) per cluster. **(G)** Dot plot of marker gene expression across clusters. Color indicates Z-score; dot size indicates the percentage of expressing cells. **(H)** Violin plots of ferroptosis suppression module scores in non-RPM populations from *Ctrl^fl/fl^* and *DKO^LysMΔ/Δ^* mice. n=3-4 animals *per* condition. Populations shown: pre-RPM mo-mac, DC, monocytes/MDM, stress-responsive monocytes, patrolling monocytes, and classical monocytes. NS, not significant (Wilcoxon rank-sum test).

**Figure S11.**
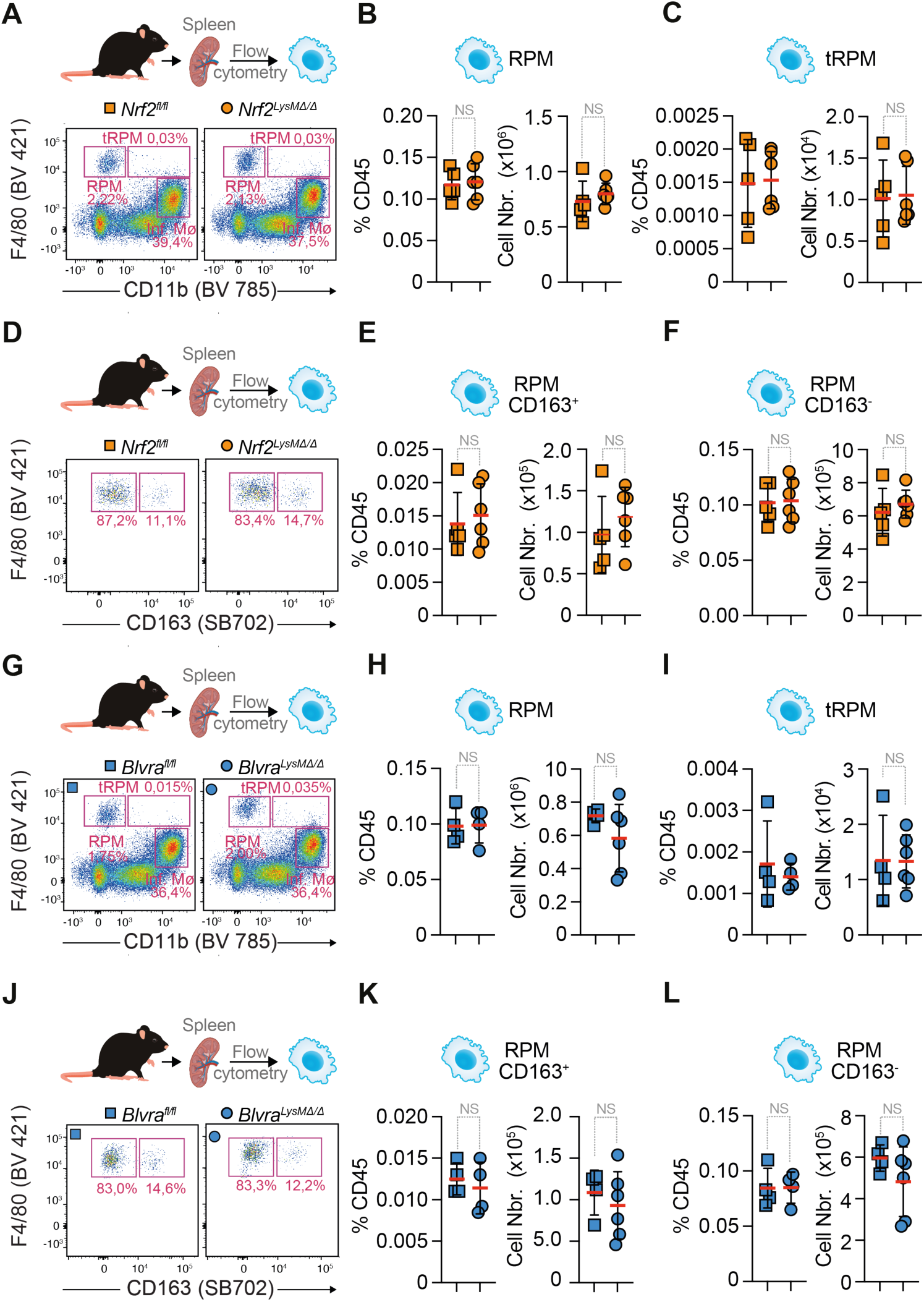
**Single deletion of NRF2 or BVRA does not alter splenic macrophage populations (#**Figure 5**). (A-C)** Representative flow cytometry plots (**A**) and quantification of RPM **(B)** and tRPM (**C**) frequency of CD45^+^ and absolute numbers in spleens from *Nrf2^fl/fl^* and *Nrf2^LysMΔ/Δ^*. **(D-F)** Representative flow cytometry plots of CD163 expression in RPM **(D)** and quantification of CD163⁺ RPM **(E)** and CD163⁻ RPM **(F)** frequency and absolute numbers in spleens from *Nrf2^fl/fl^*and *Nrf2^LysMΔ/Δ^*. **(G-I)** Representative flow cytometry plots **(G)** and quantification of RPM **(H)** and tRPM **(I)** frequency of CD45^+^ and absolute numbers in spleens from *Blvra^fl/fl^* and *Blvra^LysMΔ/Δ^*. **(J-L)** Representative flow cytometry plots of CD163 expression in RPM **(J)** and quantification of CD163⁺ RPM **(K)** and CD163⁻ RPM **(L)** frequency and absolute numbers in spleens from *Blvra^fl/fl^* and *Blvra^LysMΔ/Δ^.* Data are presented as individual data points with bar graphs showing mean ± SD. Data in (B), (C), (E), (H), (I), (K) and (I) are pooled from at least three independent experiments, n=7-11 mice *per* condition. NS, not significant, by two-tailed unpaired t-test.

**Figure S12.**
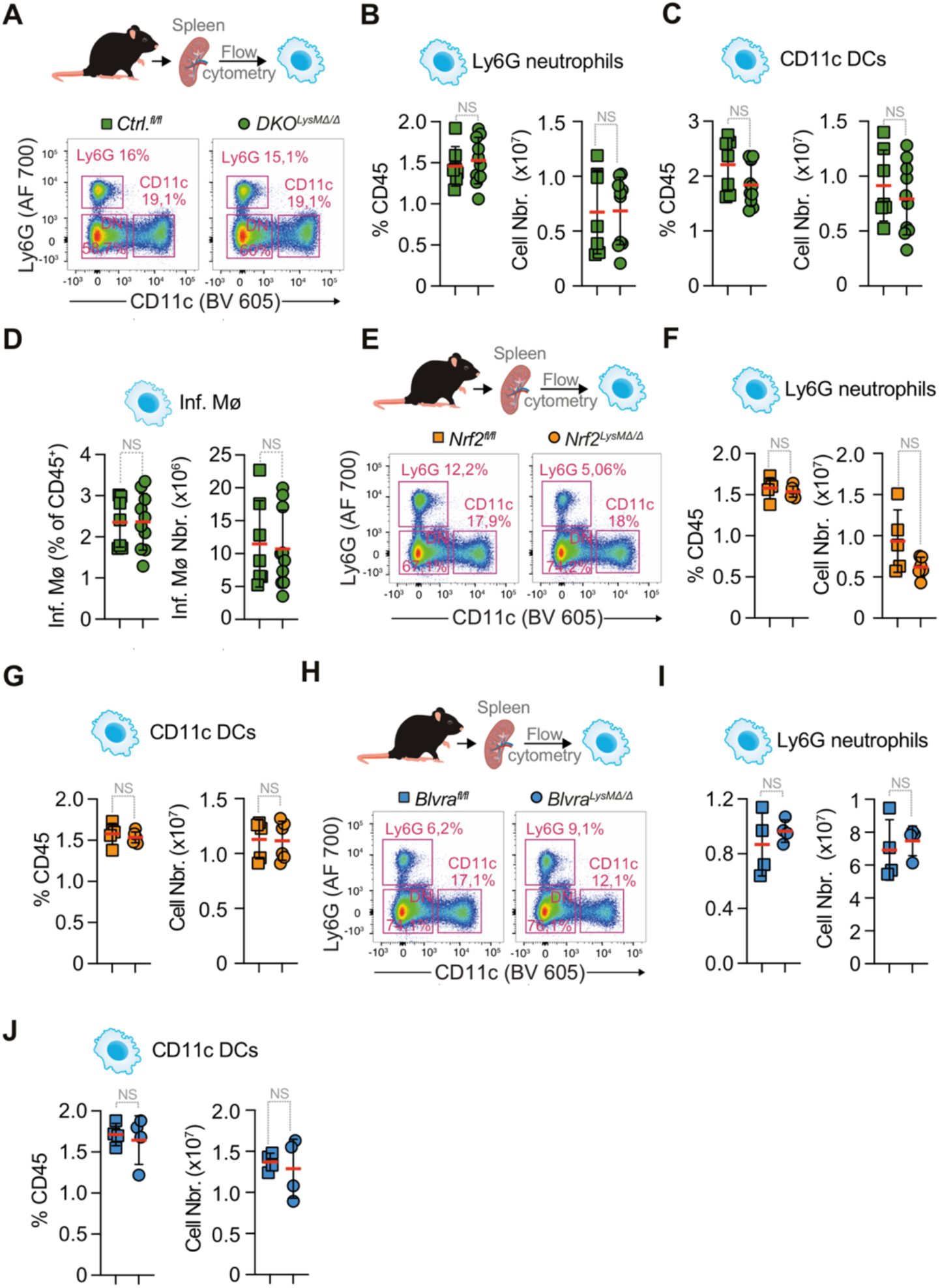
**NRF2 and BVRA deficiency does not alter splenic neutrophil, dendritic cell, or inflammatory macrophage populations (#**Figure 5**). (A–D)** Representative flow cytometry dot plots (**A**) of splenic Ly6G^+^ neutrophil and CD11c^+^ dendritic cell (DC) populations in *Ctrl^fl/fl^* and *DKO^LysMΔ/Δ^* mice, and quantification of Ly6G^+^ neutrophil (**B**), CD11c^+^ DC (**C**), and inflammatory macrophage (Inf. Mø) (**D**) frequency (% CD45) and absolute cell number. **(E-G)** Representative flow cytometry dot plots (**E**) of splenic Ly6G^+^ neutrophils and CD11c^+^ DC populations in *Nrf2^fl/fl^* and *Nrf2^LysMΔ/Δ^* mice, and quantification of Ly6G^+^ neutrophils (**F**) and CD11c^+^ DC (**G**) frequency (% CD45) and absolute cell number. **(H-J)** Representative flow cytometry dot plots (**H**) of splenic Ly6G^+^ neutrophils and CD11c^+^ DC populations in *Blvra^fl/fl^* and *Blvra^LysMΔ/Δ^*mice, and quantification of Ly6G^+^ neutrophis (**I**) and CD11c^+^ DC (**J**) frequency (% CD45) and absolute cell number. Data are presented as individual data points showing mean ± SD. Data in (B)–(D), (F,G), and (I,J) are pooled from at least three independent experiments, n=6–14 mice *per* condition. NS, not significant, by two-tailed unpaired t-test.

**Figure S13.**
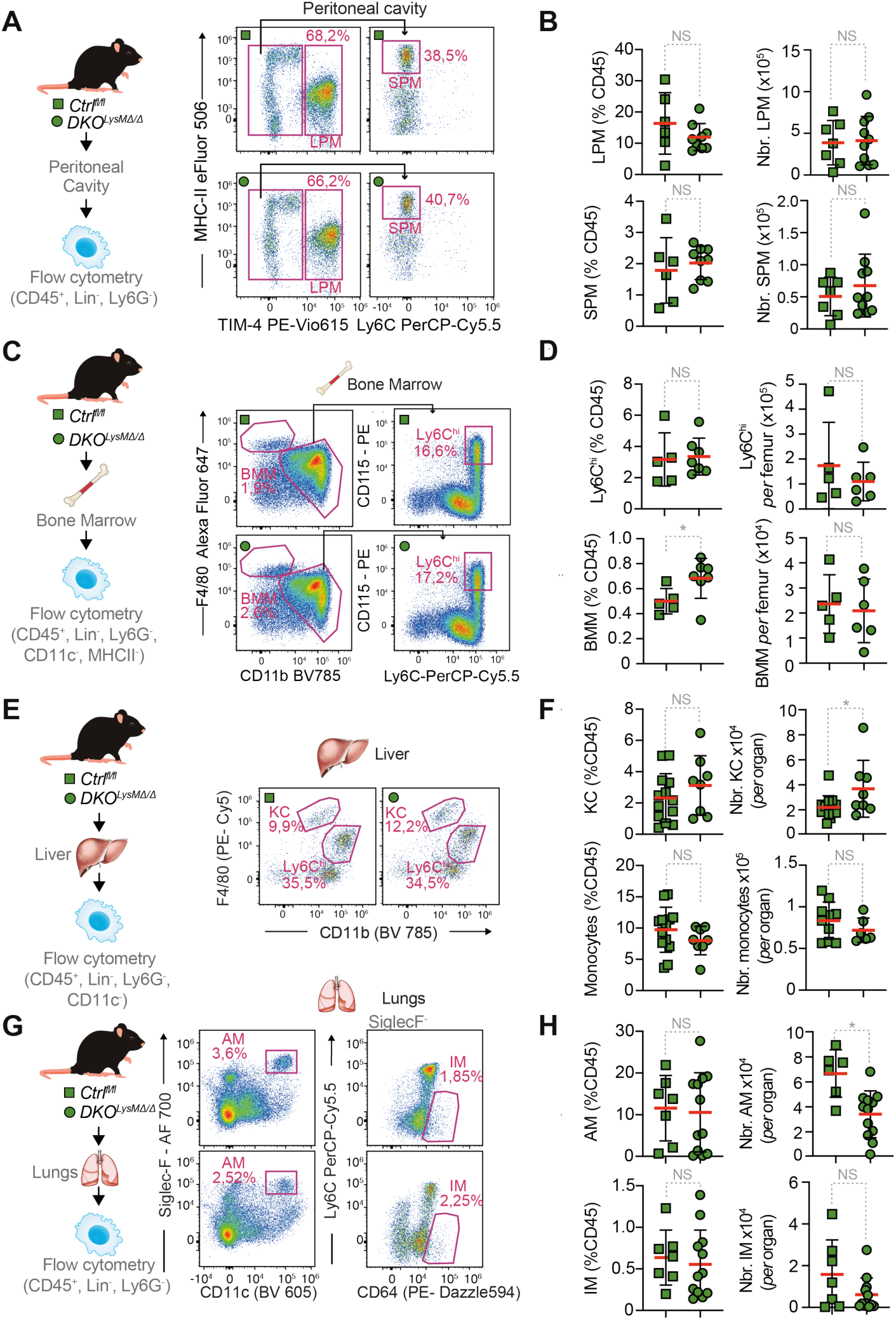
**NRF2 and BVRA deficiency does not alter tissue-resident macrophage populations across multiple organs (#**Figure 5**). (A, B)** Representative flow cytometry plots (**A**) and squantification (**B**) of large peritoneal macrophages (LPM) and small peritoneal macrophages (SPM), in *Ctrl^fl/fl^* and *DKO^LysMΔ/Δ^* mice. **(C, D)** Representative flow cytometry plots (C) and quantification (D) of bone marrow macrophage (BMM) and Ly6C^hi^ monocyte populations in *Ctrl^fl/fl^* and *DKO^LysMΔ/Δ^* mice. **(E,F)** Representative flow cytometry plots (**E**) and quantification (**F**) of hepatic myeloid populations, including Kupffer cells (KC) and hepatic monocytes, in *Ctrl^fl/fl^*and *DKO^LysMΔ/Δ^* mice. **(G, H)** Representative flow cytometry dot plots (**G**) and quantification (**H**) of lung macrophage populations, including alveolar macrophages (AM) and interstitial macrophages (IM), in *Ctrl^fl/fl^* and *DKO^LysMΔ/Δ^* mice. Data are presented as individual data points showing mean ± SD. Data in (B), (D), (F), and (H) are pooled from at least three independent experiments, n=6–14 mice *per* condition. NS, not significant; *p < 0.05; by two-tailed unpaired t-test.

**Figure S14.**
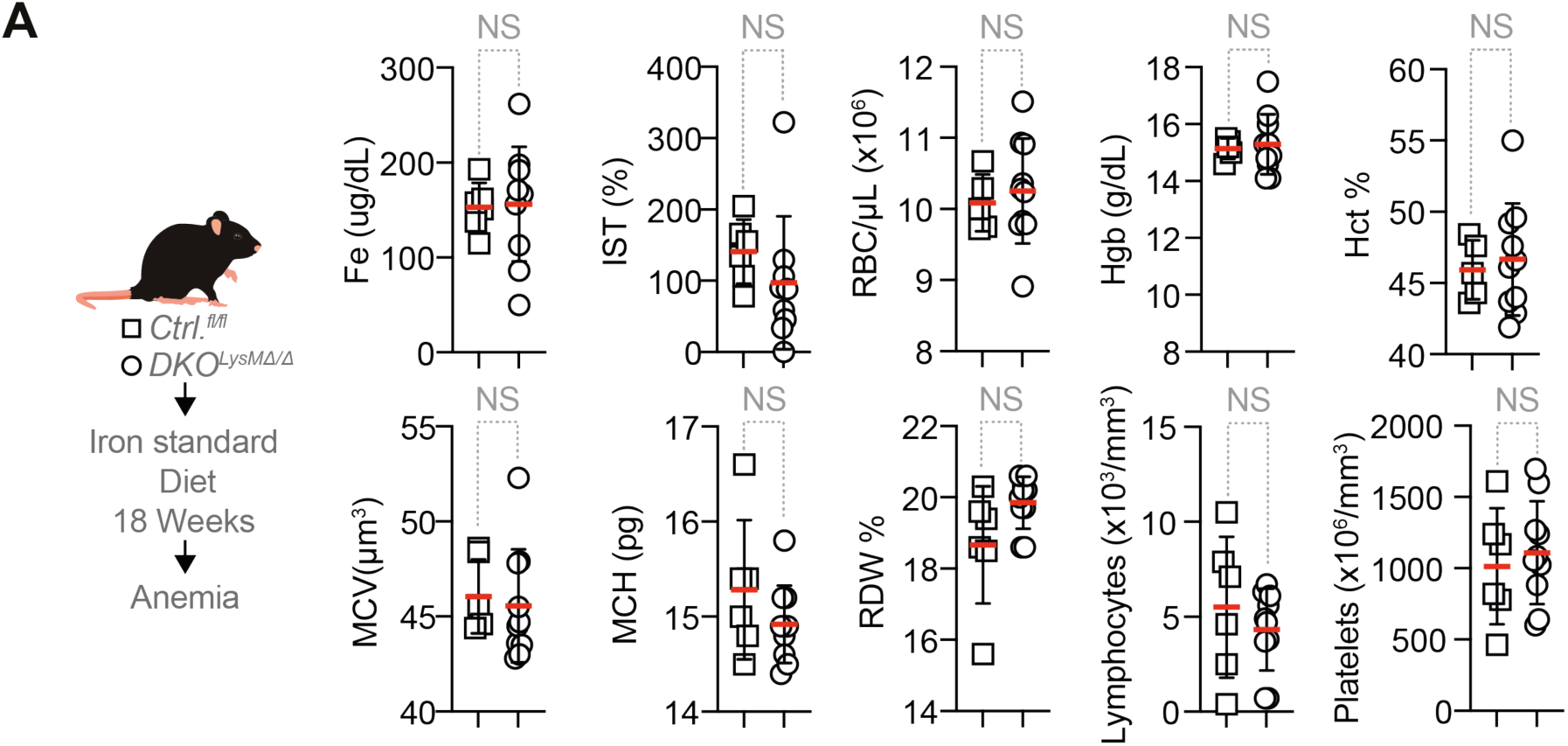
**Loss of NRF2 and BVRA expression by RPM does not induce iron-deficiency anemia at steady state (#**Figure 6**). (A)** Experimental schematic and hematological parameters in *Ctrl^fl/fl^* and *DKO^LysMΔ/Δ^* mice fed a standard diet for 18 weeks: serum iron (Fe, µg/dL), transferrin saturation index (TSI, %), red blood cell count (RBC/µL, ×10^6^), hemoglobin (Hgb, g/dL), hematocrit (Hct, %), mean corpuscular volume (MCV, µm^3^), mean corpuscular hemoglobin (MCH, pg), red cell distribution width (RDW, %), Lymphocytes (×10^3^/mm^3^), platelets (×10^6^/mm^3^). Data in **(A)** are from two independent experiments, n=6-10 mice *per* condition. NS, not significant; by two-tailed unpaired t-test or two-tailed Mann-Whitney test.

